# Rice Calcium/Calmodulin-Dependent Protein Kinase Directly Phosphorylates a Mitogen-Activated Protein Kinase Kinase to Regulate Abscisic Acid Responses

**DOI:** 10.1101/2020.10.21.348805

**Authors:** Min Chen, Lan Ni, Jing Chen, Manman Sun, Caihua Qin, Gang Zhang, Aying Zhang, Mingyi Jiang

## Abstract

Ca^2+^/calmodulin (CaM)-dependent protein kinase (CCaMK) is an important positive regulator of abscisic acid (ABA) and abiotic stress signaling in plants and is believed to act upstream of mitogen-activated protein kinase (MAPK) in ABA signaling. However, it is unclear how CCaMK activates MAPK in ABA signaling. Here, we show that OsDMI3, a rice (*Oryza sativa*) CCaMK, directly interacts with and phosphorylates OsMKK1, a MAPK kinase (MKK) in rice, in vitro and in vivo. OsDMI3 was found to directly phosphorylate Thr-25 in the N-terminus of OsMKK1, and this Thr-25 phosphorylation is OsDMI3-specific in ABA signaling. The activation of OsMKK1 and its downstream kinase OsMPK1 is dependent on Thr-25 phosphorylation of OsMKK1 in ABA signaling. Moreover, ABA treatment also induces the phosphorylation in the activation loop of OsMKK1, and the two phosphorylations in the N-terminus and in the activation loop are independent. Further analyses revealed that OsDMI3-mediated phosphorylation of OsMKK1 positively regulates ABA responses in seed germination, root growth, and tolerance to both water stress and oxidative stress. Our results indicate that OsMKK1 is a direct target of OsDMI3, and OsDMI3-mediated phosphorylation of OsMKK1 plays an important role in the activation of MAPK cascade and ABA signaling.

**One-sentence summary:** OsMKK1 is a direct target of OsDMI3, and OsDMI3-mediated phosphorylation of OsMKK1 plays an important role in the activation of MAPK cascade and ABA signaling.

The author responsible for distribution of materials integral to the findings presented in this article in accordance with the policy described in the Instructions for Authors (www.plantcell.org) is: Mingyi Jiang

## INTRODUCTION

Abscisic acid (ABA) is a major plant hormone that regulates plant growth and development and plant responses to environmental stresses. Under drought and salt stress, plants accumulate ABA, which promotes stomatal closure and triggers gene expression, thus resulting in plant adaptation to stress conditions (Cutler et al., 2010; Umezawa et al., 2010). In the ABA signaling, protein phosphorylation mediated by SNF1-related protein kinase 2s (SnRK2s), mitogen-activated protein kinases (MAPKs), calcium-dependent protein kinases (CDPKs), and calcium/calmodulin (CaM)-dependent protein kinase (CCaMK), has been shown to play an important role (Umezawa et al., 2014; Zhu, 2016a).

The MAPK cascade is one of the major signaling pathways in all eukaryotes, linking the perception of stimuli to cellular responses. The MAPK cascade is highly conserved and a typical MAPK cascade consists of three protein kinases: a MAPK kinase kinase (MAPKKK or MKKK or MEKK), a MAPK kinase (MAPKK or MKK or MEK), and a MAPK (MPK), which are activated in a sequential phosphorylation manner. Once activated, MAPKs phosphorylate various downstream protein targets, such as transcription factors, phospholipases, protein kinases, metabolic enzymes, cytoskeletal and microtubule-associated proteins, leading to the activation of cellular responses (Umezawa et al., 2014; de Zelicourt et al., 2016; Dóczi and Bögre, 2018; Zhang et al., 2018). There are 80 MAPKKKs, 10 MAPKKs and 20 MAPKs in Arabidopsis (MAPK Group, 2002; Colcombet and Hirt, 2008), and 75 MAPKKKs, 8 MAPKKs and 17 MAPKs in rice (Hamel et al., 2006; Rao et al., 2010). Several complete MAPK cascades have been identified to be involved in plant response to biotic and abiotic stresses. In Arabidopsis, diverse pattern recognition receptors (PRRs) activate two MAPK cascades: the first one is composed of MEKK1-MKK1/2-MPK4 (Suarez-Rodriguez et al., 2007; Gao et al., 2008), the second one is composed of MAPKKK3/5-MKK4/5-MPK3/6 (Bi et al., 2018; Sun et al., 2018). Similarly, rice OsMAPKKK11/18-OsMKK4-OsMPK3/6 (Yamada et al., 2017) and OsMAPKKK24-OsMKK4-OsMPK3/6 (Wang et al., 2017) were also reported to function in chitin signaling. In the response of plants to abiotic stresses, a recent study revealed that the MEKK1-MKK2-MPK4 cascade positively regulates freezing tolerance, and the MKK4/5-MPK3/6 cascade negatively regulates freezing tolerance (Zhao et al., 2017). In ABA signaling, it was shown that the ABA-activated MAPKKK17/18-MKK3-MPK1/2/7/14 cascade is involved in stress signaling (Danquah et al., 2015) and timing of senescence (Matsuoka et al., 2015). A recent study reported that the MAPKKK20 (AIK1)-MKK5-MPK6 cascade functions in the ABA regulation of primary root growth and stomatal response (Li et al., 2017).

CCaMK, as a decoder for Ca^2+^ signals, has been shown to be a crucial regulator of root nodule and arbuscular mycorrhizal symbioses (Singh and Parniske, 2012; Poovaiah et al., 2013). CCaMK has been also shown to be involved in the responses of plants to abiotic stresses (Ma et al., 2012; Shi et al., 2012, 2014; Zhu et al., 2016b; Ni et al., 2019) and biotic stresses (Wang et al., 2015). Previous studies have indicated that the rice CCaMK OsDMI3 is a positive regulator of ABA responses, including seed germination, root growth, antioxidant defense, and tolerance to both water stress and oxidative stress (Shi et al., 2012, 2014; Ni et al., 2019). A recent study revealed that in the absence of ABA, the type 2C protein phosphatase (PP2C) OsPP45 directly interacts with OsDMI3 to inactivate OsDMI3 by dephosphorylating, and in the presence of ABA, ABA-induced H_2_O_2_ production inactivates OsPP45, resulting in OsDMI3 activation (Ni et al., 2019). OsDMI3 was shown to function upstream of OsMPK1 (also called OsMPK6), a major ABA-activated MAPK, to regulate the antioxidant defense systems in ABA signaling (Shi et al., 2014). However, the molecular mechanism of OsDMI3-mediated activation of OsMPK1 in ABA signaling remains to be determined. Here, we show that OsDMI3 directly interacts with and phosphorylates the upstream activator of OsMPK1, OsMKK1 in vitro and in vivo, and OsDMI3-mediated phosphorylation of OsMKK1 is required for the activation of OsMKK1-OsMPK1 cascade in ABA signaling. Genetic evidence demonstrates that OsDMI3-mediated phosphorylation of OsMKK1 plays an important role in ABA signaling. These findings uncover an important non-canonical MKK activation mechanism, which directly connects CCaMK to the MAPK cascade in ABA signaling.

## RESULTS

### OsDMI3 Does Not Interact Directly with OsMPK1, But Interacts with Its Upstream Activators OsMKK1 and OsMKK6

To determine whether there exists a direct interaction between OsDMI3 and OsMPK1 in plant cells, yeast two-hybrid (Y2H) assays and bimolecular fluorescence complementation (BiFC) assays were performed. Experimental results showed that OsDMI3 did not interact with OsMPK1 either in yeast cells (Supplemental Figure 1A) or in onion epidermis cells (Supplemental Figure 1B). Moreover, an in vitro phosphorylation assay showed that OsDMI3 did not directly phosphorylate OsMPK1 (Supplemental Figure 1C). These results suggest that OsDMI3 does not directly activate OsMPK1 in plant cells.

In a canonical MAPK cascade, MAPK is activated by its upstream MKK. In rice, OsMPK1 was shown to interact with OsMKK1 (OsMEK2), OsMKK6 (OsMEK1), OsMKK3 (OsMEK8a), OsMKK4 (OsMEK6), OsMKK5 (OsMEK7b), and OsMKK10-2 (OsMEK3) (Singh et al. 2012). To determine whether OsDMI3 can interact with these MKKs, Y2H and BiFC assays were conducted. OsDMI3 was shown to interact with the group A MKKs, OsMKK1 and OsMKK6, in both yeast cells (Figure 1A; Supplemental Figure 2A) and onion epidermis cells (Figure 1C; Supplemental Figure 2B). Further, glutathione S-transferase (GST) pull-down assays showed that OsDMI3 was able to directly interact with OsMKK1 and OsMKK6 in vitro (Figure 1B). The OsDMI3-OsMKK1 and OsDMI3-OsMKK6 interactions were also detected in co-immunoprecipitation (Co-IP) assays when OsDMI3-Myc was co-expressed with OsMKK1-Flag or OsMKK6-Flag in rice protoplasts. Immunoblot (IB) analyses using an anti-Flag antibody revealed that both OsMKK1-Flag and OsMKK6-Flag interacted with OsDMI3-Myc in rice protoplasts (Figure 1D).

**Figure 1.**
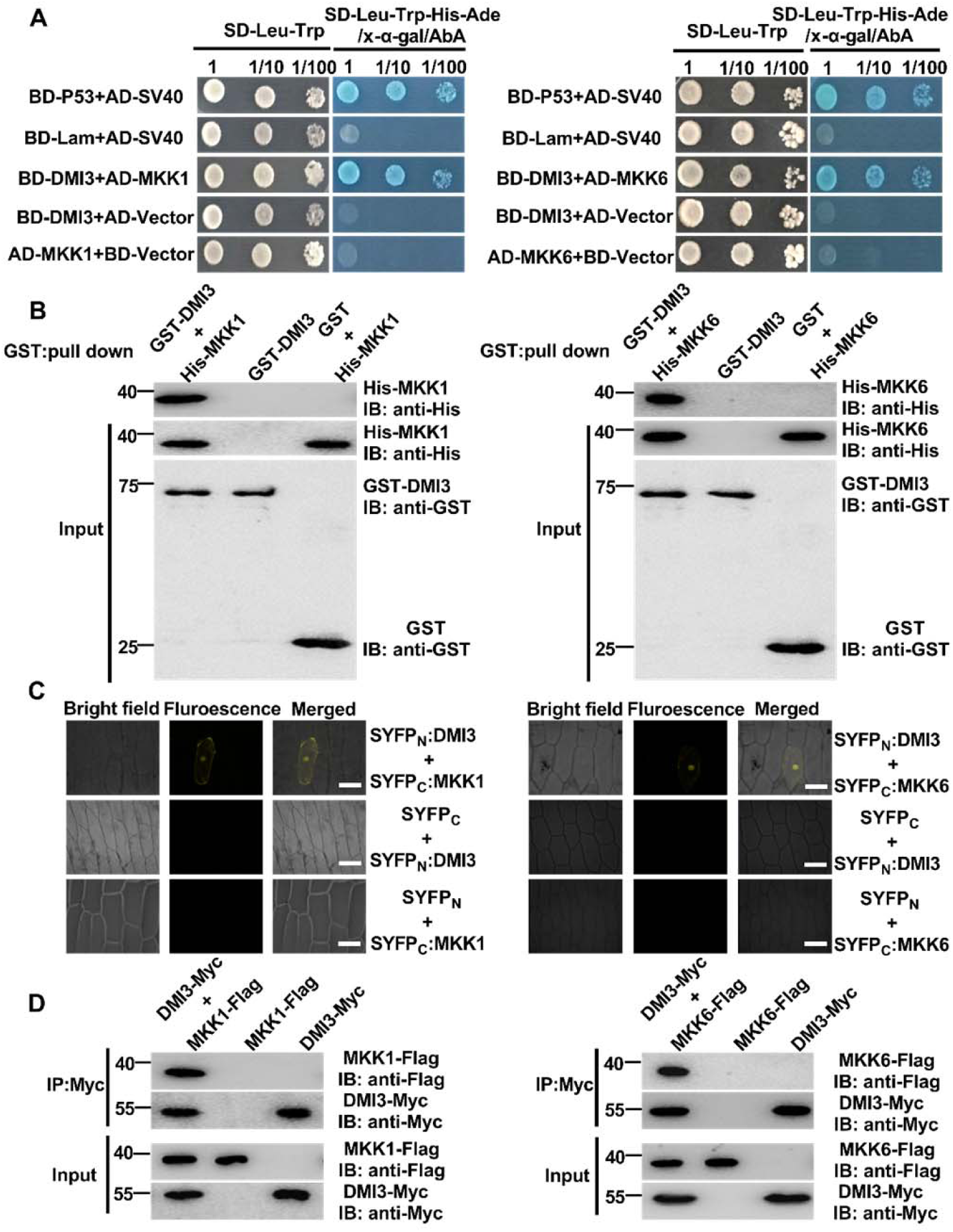
OsDMI3 Interacts with OsMKK1/OsMKK6 Both In Vitro and In Vivo. **(A)** Y2H assay. SD-Trp-Leu-His-Ade/AbA/X-α-gal medium was used for testing the interaction between OsDMI3 and OsMKK1 (left) or OsMKK6 (right). BD-P53/AD-SV40 was used as a positive control, and BD-Lam/AD-SV40 was used as a negative control. **(B)** GST pull-down assay. The equal amount of GST and GST-DMI3 were incubated with His-OsMKK1 (left) or His-OsMKK6 (right) in GST beads. GST-OsDMI3 was detected with anti-GST antibody, and both His-OsMKK1 and His-OsMKK6 were detected with anti-His antibody. **(C)** BiFC analysis. The indicated constructs were transiently expressed in onion epidermis cells. Co-expression of *SYFP*_*N*_*-OsDMI3* plus *SYFP*_*C*_ or *SYFP*_*C*_*-OsMKK1* plus *SYFP*_*N*_ (left) or *SYFP*_*C*_*-OsMKK6* plus *SYFP*_*N*_ (right) were used as negative controls. Scale bars, 100 µm. **(D)** Co-IP analysis. OsDMI3-Myc and OsMKK1-Flag (left) or OsMKK6-Flag (right) were co-transformed into rice protoplasts, and the protein extracts were immunoprecipitated (IP) using an anti-Myc antibody and were detected with anti-Flag (OsMKK1/OsMKK6) and anti-Myc (OsDMI3) antibodies. Protein input was shown by IB analysis of protein extracts before immunoprecipitation and antibodies against the respective tags. Molecular mass markers in kD were shown on the left. All experiments were repeated at least three times with similar results.

To determine which regions of both OsDMI3 and OsMKK1 are required for the interaction, a series of deletion derivatives of the two proteins were generated and then tested for the interaction using Y2H assay and luciferase complementation imaging (LCI) assay. Y2H assays showed that OsMKK1 interacted with the EF (helix-loop-helix) hands domain (337 to 516 amino acids) of OsDMI3 (Supplemental Figure 3A). LCI assays confirmed the in vivo interaction between OsMKK1 and the EF hands domain of OsDMI3 (Supplemental Figure 3B). Similarly, OsDMI3 was shown to interact with the N-terminus domain (1 to 65 amino acids) of OsMKK1 in Y2H (Supplemental Figure 4A) and LCI assays (Supplemental Figure 4B). Moreover, the interaction domains between OsDMI3 and OsMKK6 were also tested by Y2H assay and by LCI assay. Experimental results showed that the EF hands domain (337 to 516 amino acids) in the C-terminus of OsDMI3 (Supplemental Figures 5A and 5B) interacted with the N-terminus domain (1 to 71 amino acids) of OsMKK6 (Supplemental Figures 6A and 6B). Taken together, these results indicate that the OsDMI3-OsMKK1/OsMKK6 interaction involves the EF hands domain of OsDMI3 and the N-terminus domain of OsMKK1/OsMKK6.

### OsDMI3-Dependent Activation of OsMKK1 But Not OsMKK6 in ABA Signaling

To determine whether OsDMI3 mediates the activation of both OsMKK1 and OsMKK6 in ABA signaling, we first tested if ABA induces the activation of both OsMKK1 and OsMKK6 in planta. Two independent *OsMKK1*- or *OsMKK6*-overexpressing (OE) lines (*OsMKK1*-OE1, *OsMKK1*-OE2, *OsMKK6*-OE1, and *OsMKK6*-OE2) and two independent *OsMKK1*- or *OsMKK6*-knockout (KO) lines (*osmkk1*-KO1, *osmkk1*-KO2, *osmkk6*-KO1, and *osmkk6*-KO2) were generated, and then the specific anti-OsMKK1 and anti-OsMKK6 antibodies were prepared. The activities of OsMKK1 and OsMKK6 in rice plants were determined by an immunoprecipitation kinase assay with myelin basic protein (MBP) as a substrate. In the *osmkk1*-KO lines (Supplemental Figure 7A) and the *osmkk6*-KO lines (Supplemental Figure 7D) generated by CRISPR/Cas9 system, the activities of both OsMKK1 (Supplemental Figures 7B and 7C) and OsMKK6 (Supplemental Figures 7E and 7F) were undetectable. By contrast, the *OsMKK1*-OE lines and the *OsMKK6*-OE lines exhibited high activities of OsMKK1 and OsMKK6. ABA treatment induced a rapid increase in the activities of both OsMKK1 (Figures 2A and 2B) and OsMKK6 (Figures 2C and 2D) in rice leaves, with a significant increase at 30 min of ABA treatment and a maximal increase at 90 min of ABA treatment. The specificity of the anti-OsMKK1 and anti-OsMKK6 antibodies was proven by immunoblotting analysis using the proteins extracted from the leaves of wild type (WT), *OsMKK1*-OE1, and *osmkk1*-KO1 plants (Supplemental Figure 8A) or *OsMKK6*-OE1 and *osmkk6*-KO1 plants (Supplemental Figure 8B).

**Figure 2.**
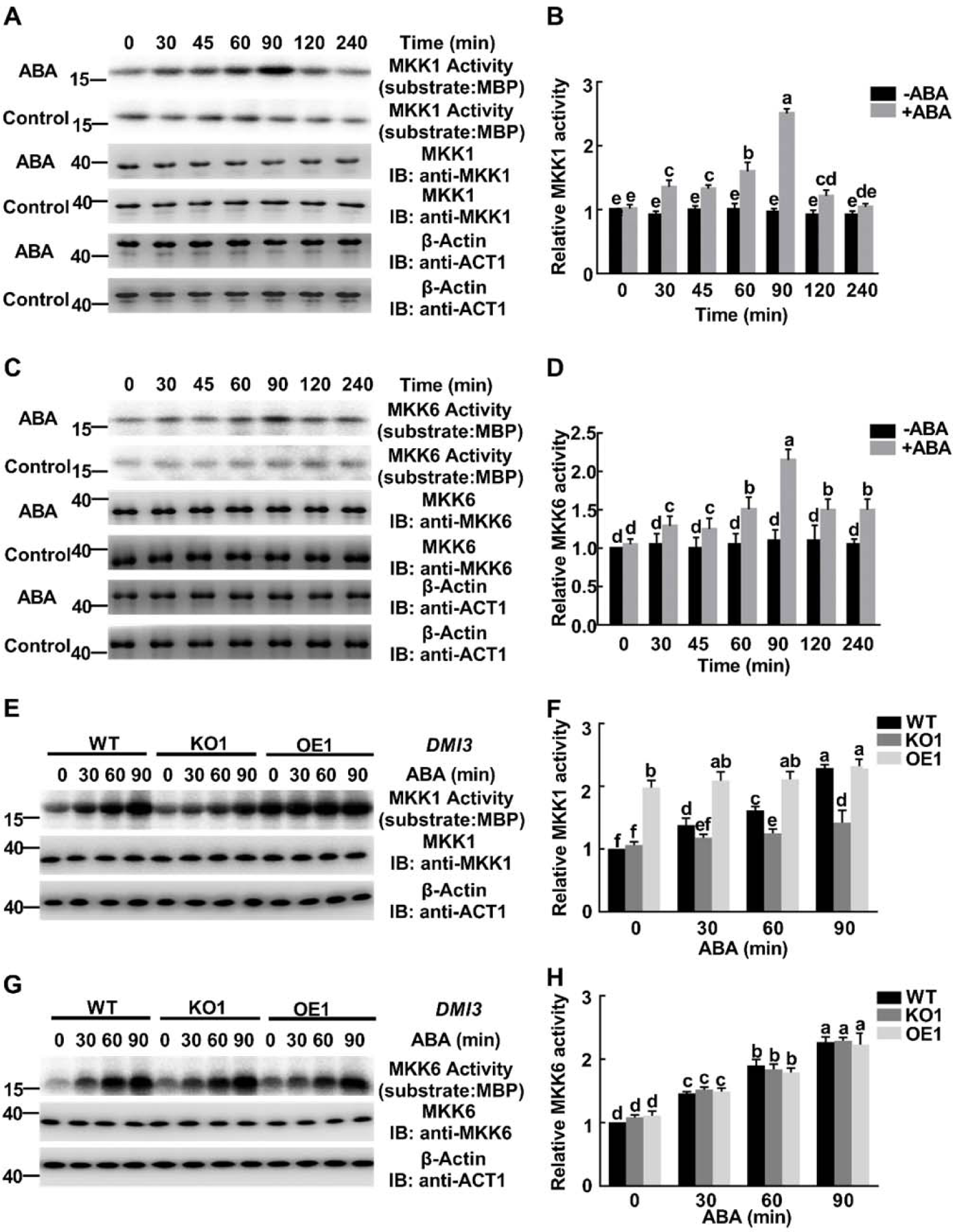
The Activation of OsMKK1 But Not OsMKK6 Is OsDMI3-Dependent in ABA Signaling. **(A)** and **(C)** ABA-induced increases in the activities of OsMKK1 **(A)** and OsMKK6 **(C)** in rice leaves. The rice seedlings were treated with 100 µM ABA for various times as indicated. OsMKK1 and OsMKK6 were immunoprecipitated from rice leaves, and the activities of OsMKK1 and OsMKK6 were analyzed by immunoprecipitation kinase assay using MBP as substrate. OsMKK1 and OsMKK6 input were analyzed by IB using anti-OsMKK1 antibody and anti-OsMKK6 antibody, respectively. β-actin was used as the total protein loading control. Molecular mass markers in kD were shown on the left. **(B)** and **(D)** The relative activities of OsMKK1 **(B)** in **(A)** and OsMKK6 **(D)** in **(C)**. Kinase activities were quantitated by ImageJ software, and the activities of OsMKK1 and OsMKK6 in the control leaves treated with ABA for 0 min were set to 1, respectively. **(E)** and **(G)** The activities of OsMKK1 **(E)** and OsMKK6 **(G)** in *OsDMI3*-OE1, *osdmi3*-KO1 and WT. The rice seedlings were treated with 100 µM ABA for 30, 60, and 90 min, and the activities of OsMKK1 and OsMKK6 were analyzed by immunoprecipitation kinase assay using MBP as substrate. OsMKK1 and OsMKK6 input were analyzed by IB using anti-OsMKK1 antibody and anti-OsMKK6 antibody, respectively. β-actin was used as the total protein loading control. Molecular mass markers in kD were shown on the left. **(F)** and **(H)** The relative activities of OsMKK1 **(F)** in **(E)** and OsMKK6 **(H)** in **(G)**. Kinase activities were quantitated by ImageJ software, and the activities of OsMKK1 and OsMKK6 in WT in the control were set to 1, respectively. All experiments were repeated at least three times with similar results. In **(B) (D) (F)** and **(H)**, values are means ± SEM of three independent experiments. Means denoted by the same letter did not significantly differ at P < 0.05 according to Duncan’s multiple range test.

Then, we used the *OsDMI3*-knockout mutant *osdmi3*-KO1 and the *OsDMI3*-overexpressing line *OsDMI3*-OE1 to investigate whether OsDMI3 is required for the ABA-induced activation of both OsMKK1 and OsMKK6. Under the nontreated conditions, the *OsDMI3*-OE1 plants showed increased activity of OsMKK1, but the *osdmi3*-KO1 plants showed no obvious change in OsMKK1 activity, compared with WT plants (Figures 2E and 2F). After ABA treatment, the ABA-induced increase in OsMKK1 activity was slightly enhanced in *OsDMI3*-OE1 plants, but was substantially inhibited in *osdmi3*-KO1 plants, indicating that ABA-induced activation of OsMKK1 is OsDMI3-dependent (Figures 2E and 2F). However, there was no difference among *OsDMI3*-OE1, *osdmi3*-KO1 and WT plants in the activity of OsMKK6 under the nontreated conditions, and ABA treatment induced the same increase in OsMKK6 activity in these plants, indicating that ABA-induced activation of OsMKK6 is OsDMI3-independent in rice plants (Figures 2G and 2H).

Finally, we investigated whether ABA-induced activation of both OsMKK1 and OsMKK6 is calcium-dependent. Treatment with 2 mM CaCl_2_ induced the activation of OsMKK1 (Supplemental Figures 9A and 9B) but not OsMKK6 (Supplemental Figures 10A and 10B), and pretreatments with the Ca^2+^ chelator ethyleneglycoltetraacetic acid (EGTA) and the Ca^2+^ channel blocker LaCl_3_ inhibited ABA-induced increase in OsMKK1 activity (Supplemental Figures 9C and 9D) but not in OsMKK6 activity (Supplemental Figures 10C and 10D), indicating that ABA-induced activation of OsMKK1 is Ca^2+^-dependent, and ABA-induced activation of OsMKK6 is Ca^2+^-independent. Furthermore, Ca^2+^-induced increase in OsMKK1 activity was greatly reduced in *osdmi3*-KO1 plants, indicating that OsDMI3 makes a major contribution to Ca^2+^-induced activation of OsMKK1 in rice plants (Supplemental Figures 9E and 9F).

On the other hand, we also investigated whether OsMKK1 mediates the activation of OsDMI3 in ABA signaling. Under the nontreated conditions, there was no difference among *OsMKK1*-OE1, *osmkk1*-KO1 and WT plants in the activity of OsDMI3, and ABA treatment induced the same increase in OsDMI3 activity in these plants, indicating that OsMKK1 is not involved in the activation of OsDMI3 in ABA signaling (Supplemental Figures 11A and 11B).

### OsDMI3 Phosphorylates OsMKK1 But Not OsMKK6 in ABA Signaling

To determine if OsMKK1 or OsMKK6 can be phosphorylated by OsDMI3, both in vitro and in vivo protein phosphorylation assays were conducted. In vitro kinase assays showed that OsDMI3 extracted from ABA-treated rice leaves phosphorylated both His-OsMKK1 (Figure 3A) and His-OsMKK6 (Figure 3B). To investigate whether this phosphorylation by OsDMI3 also occurs in plant cells, *OsMKK1* or *OsMKK6* were introduced into the protoplasts of WT and *osdmi3*-KO1, and the transfected protoplasts were treated with ABA. The phosphorylated proteins of OsMKK1 and OsMKK6 were detected by immunoblotting using biotinylated Phos-tag. Under the nontreated conditions, there was no difference between WT and *osdmi3*-KO1 protoplasts with transiently expressed *OsMKK1* in the phosphorylation level of OsMKK1 (Figures 3C and 3D), and also no difference between WT and *osdmi3*-KO1 protoplasts with transiently expressed *OsMKK6* in the phosphorylation level of OsMKK6 (Figures 3E and 3F). ABA treatment induced a significant increase in the phosphorylation level of OsMKK1 in the WT protoplasts, with a maximal increase at 10 min of ABA treatment, but the ABA-induced increase was substantially inhibited in the *osdmi3*-KO1 protoplasts, indicating that ABA-induced OsMKK1 phosphorylation is OsDMI3-dependent (Figures 3C and 3D). However, although ABA also induced an increase in OsMKK6 phosphorylation in the WT protoplasts, this increase was not affected in the *osdmi3*-KO1 protoplasts, indicating that ABA-induced OsMKK6 phosphorylation is OsDMI3-independent (Figures 3E and 3F).

**Figure 3.**
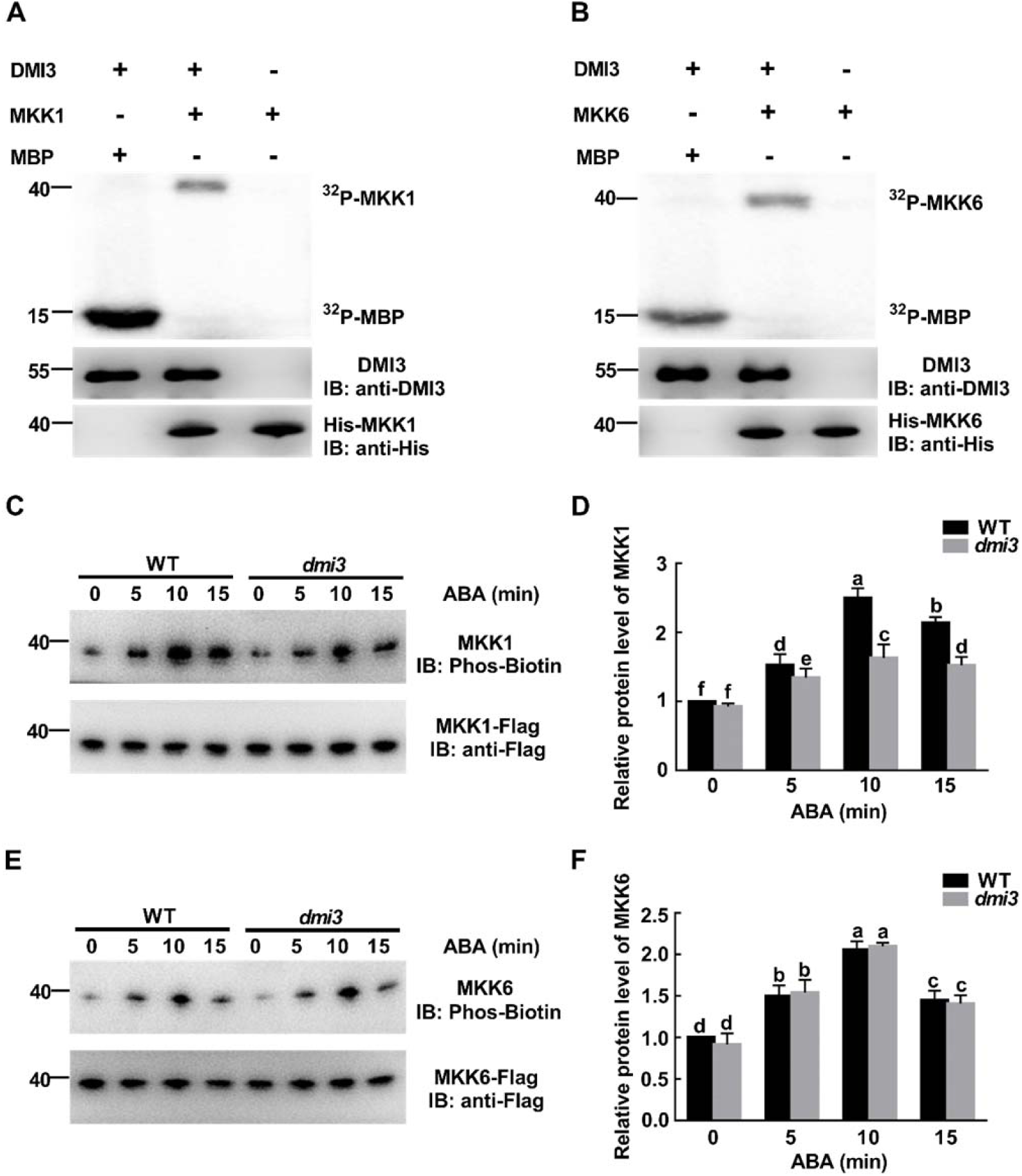
OsDMI3 Phosphorylates OsMKK1 But Not OsMKK6 in ABA Signaling. **(A)** and **(B)** OsDMI3 phosphorylates both OsMKK1 **(A)** and OsMKK6 **(B)** in vitro. Protein extracts from ABA-treated rice leaves were immunoprecipitated with anti-OsDMI3 antibody. His-OsMKK1, His-OsMKK6 and MBP were used as substrates and subjected to an in-gel kinase assay (top). The amount of OsDMI3 protein was determined by IB with anti-OsDMI3 antibody (middle). The recombinant proteins of His-OsMKK1 and His-OsMKK6 were determined by IB with anti-His antibody (bottom). **(C)** and **(E)** OsDMI3 phosphorylates OsMKK1 **(C)** but not OsMKK6 **(E)** in ABA signaling. *OsMKK1-Flag* or *OsMKK6-Flag* were introduced into rice protoplasts of WT and *osdmi3*-KO1, and the transfected protoplasts were treated with 10 µM ABA for 5, 10, and 15 min. Protein extracts were immunoprecipitated with anti-Flag antibody and were detected by IB with anti-Flag antibody (bottom). The phosphorylated OsMKK1 or OsMKK6 proteins were detected by IB using biotinylated Phos-tag (top). **(D)** and **(F)** Relative protein levels of phosphorylated OsMKK1 **(D)** and OsMKK6 **(F)** quantitated by ImageJ software. The protein levels of phosphorylated OsMKK1 and OsMKK6 in WT in the control were set to 1, respectively. All experiments were repeated at least three times with similar results. In **(D)** and **(F)**, values are means ± SEM of three independent experiments. Means denoted by the same letter did not significantly differ at P < 0.05 according to Duncan’s multiple range test. Molecular mass markers in kD were shown on the left.

### OsDMI3 Phosphorylates Thr-25 in the N-Terminus of OsMKK1

We next sought to determine which amino acid residues in OsMKK1 are phosphorylated by OsDMI3. The recombinant OsMKK1 was pretreated with calf intestine alkaline phosphatase (CIAP) to remove the pre-existing phosphorylation events and then incubated with OsDMI3, and the phosphorylation sites were determined by liquid chromatography-tandem mass spectrometry (LC-MS/MS) analysis. Thr-25 of OsMKK1 was identified as the residue phosphorylated by OsDMI3 (Supplemental Table 1 and Supplemental Figure 12). To further determine the phosphorylation site of OsMKK1 by OsDMI3, Thr-25 of OsMKK1 was mutated to Ala to create the non-phosphorylatable mutant OsMKK1^T25A^. In vitro kinase assays showed that the OsMKK1^T25A^ mutant protein was not phosphorylated by OsDMI3 (Figure 4A), indicating that Thr-25 in OsMKK1 is the residue phosphorylated by OsDMI3 in vitro.

**Figure 4.**
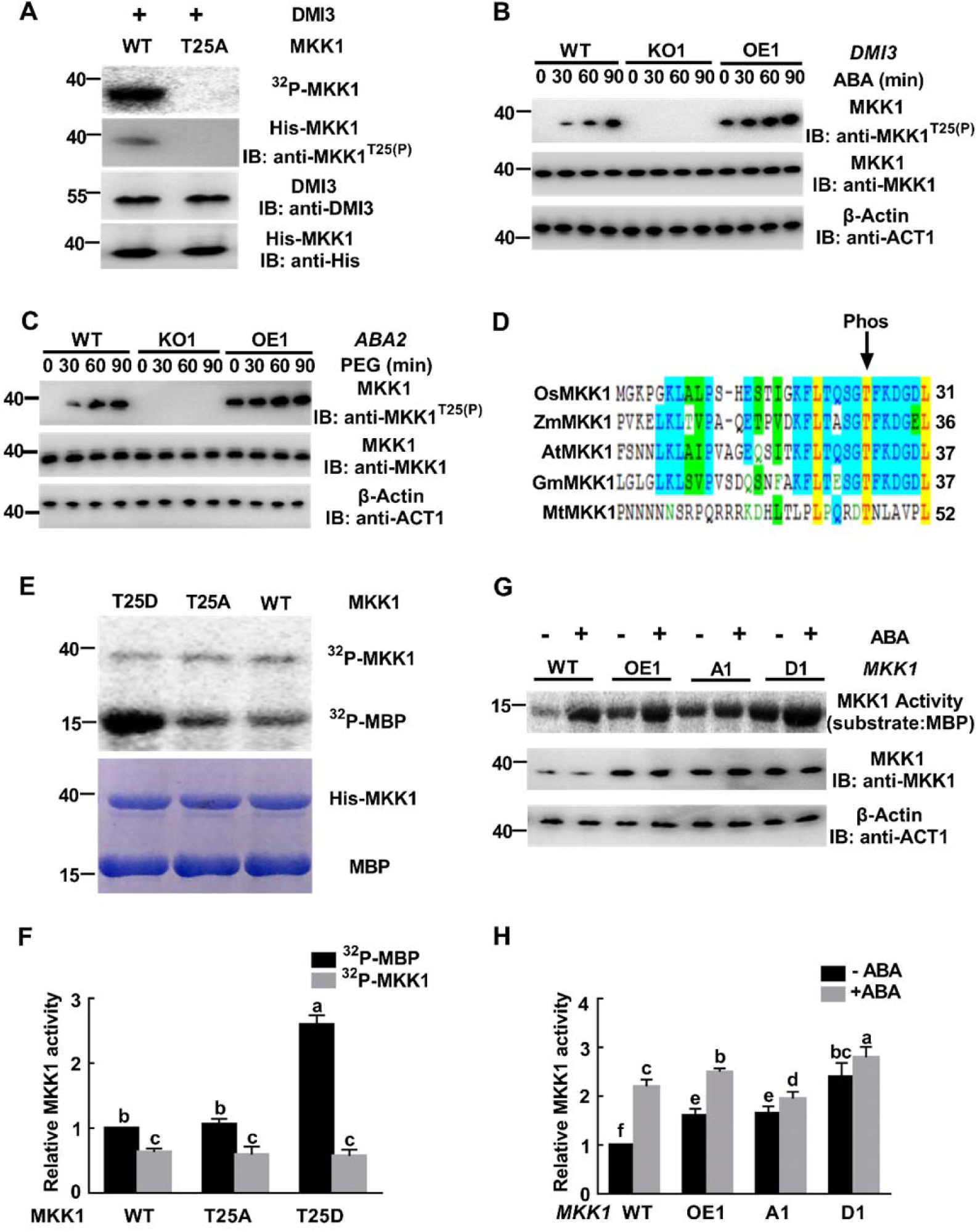
OsDMI3 Directly Phosphorylates OsMKK1 at Thr-25 to Regulate OsMKK1 Activity. **(A)** In vitro phosphorylation of OsMKK1 and mutated OsMKK1 (T25A) by OsDMI3. Protein extracts from ABA-treated rice leaves were immunoprecipitated with anti-OsDMI3 antibody. His-OsMKK1 and three mutated His-OsMKK1 proteins were used as substrates and subjected to an in-gel kinase assay. Thr-25 phosphorylation was analyzed by IB using anti-pT25 OsMKK1 antibody. OsDMI3 protein was tested by IB with anti-OsDMI3 antibody, and His-OsMKK1 and mutated His-OsMKK1 proteins were determined by IB with anti-His antibody. **(B)** Thr-25 phosphorylation is OsDMI3-dependent in ABA signaling. *OsDMI3*-OE1, *osdmi3*-KO1 and WT plants were treated with 100 µM ABA for 30, 60, and 90 min, and Thr-25 phosphorylation was tested by IB with anti-pT25 OsMKK1 antibody. OsMKK1 input was analyzed by IB with anti-OsMKK1 antibody. β-actin was used as total protein loading control. **(C)** ABA-dependence of Thr-25 phosphorylation under water stress. *OsABA2*-OE1, *osaba2*-KO1 and WT plants were treated with 20% PEG 4000 for 30, 60, and 90 min, and Thr-25 phosphorylation was tested by IB with anti-pT25 OsMKK1 antibody. OsMKK1 input was analyzed by IB with anti-OsMKK1 antibody. β-actin was used as total protein loading control. **(D)** The N-terminal Thr-25 residue of OsMKK1 is conserved in monocots and dicots. N-terminal sequences of MKKs from *Zea mays* (ZmMKK1), *Arabidopsis thaliana* (AtMKK1), *Glycine max* (GmMKK1), *Medicago truncatula* (MtMKK1) and *Oryza sativa* (OsMKK1) were aligned. The number at the end of each line indicates the coordinate of the last residue. **(E)** OsMKK1^T25D^ enhances the activity of OsMKK1 in vitro. The upper gel shows substrate phosphorylation (MBP as substrate) and autophosphorylation activity of OsMKK1, OsMKK1^T25A^ and OsMKK1^T25D^, and the bottom gel shows the corresponding Coomassie staining. **(F)** The relative activities of substrate phosphorylation and autophosphorylation in **(E)**. The activities of autophosphorylation and substrate phosphorylation of OsMKK1 were quantitated by ImageJ software, and the substrate phosphorylation activity of OsMKK1 was set to 1. **(G)** Thr-25 phosphorylation is required for the activation of OsMKK1 in ABA signaling. *OsMKK1*-OE1, *OsMKK1*^*T25A*^-A1, *OsMKK1*^*T25D*^-D1, and WT plants were treated with 100 µM ABA for 90 min, and the activity of OsMKK1 was analyzed by immunoprecipitation kinase assay using MBP as substrate. OsMKK1 input was analyzed by IB with anti-OsMKK1 antibody. β-actin was used as the total protein loading control. **(H)** The relative activity of OsMKK1 in **(G)**. Kinase activity was quantitated by ImageJ software. The activity of OsMKK1 in untreated WT was set to 1. All experiments were repeated at least three times with similar results. In **(F)** and **(H)**, values are means ± SEM of three independent experiments. Means denoted by the same letter did not significantly differ at P < 0.05 according to Duncan’s multiple range test. Molecular mass markers in kD were shown on the left.

To determine if OsDMI3 is responsible for the phosphorylation of Thr-25 in OsMKK1 in rice plants, a specific anti-phospho-Thr-25 antibody was prepared. This antibody specifically recognized the phosphorylated Thr-25 of OsMKK1, but not the two mutants of OsMKK1 in which Thr-25 was converted to either Ala (OsMKK1^T25A^) or Asp (OsMKK1^T25D^, a phosphomimetic mutant) (Supplemental Figures 13A to 13C). Under the nontreated conditions, OsMKK1 Thr-25 phosphorylation was not observed in WT and *osdmi3*-KO1 plants, but was strong in *OsDMI3*-OE1 plants (Figure 4B). ABA treatment induced a progressive increase in OsMKK1 Thr-25 phosphorylation in WT plants in a time-dependent manner during the 90-min treatment, and further enhanced OsMKK1 Thr-25 phosphorylation in *OsDMI3*-OE1 plants. However, this phosphorylation was completely blocked in *osdmi3*-KO1 plants, indicating that OsMKK1 Thr-25 phosphorylation is specifically dependent on OsDMI3 in ABA signaling (Figure 4B).

To further determine whether endogenous ABA also plays such a role under water stress, the *OsABA2*-overexpressing line (*OsABA2*-OE1) and the *OsABA2*-knockout line (*osaba2*-KO1) were generated (Supplemental Figures 14A and 14B). It has been shown that ABA2 can catalyze the conversion of xanthoxin to abscisic aldehyde in ABA biosynthesis (Cheng et al., 2002; González-Guzmán et al., 2002; Ma et al., 2016). In *osaba2*-KO1 plants, the content of ABA was about 20% of WT plants, but in *OsABA2*-OE1 plants, the content of ABA was about 2.6 times that of WT plants (Supplemental Figure 14B). Water stress induced by polyethylene glycol (PEG) led to a progressive increase in OsMKK1 Thr-25 phosphorylation in WT plants in a time-dependent manner during the 90-min treatment, and the increase was further enhanced in *OsABA2*-OE1 plants (Figure 4C). By contrast, this phosphorylation induced by water stress was completely blocked in *osaba2*-KO1 plants, indicating that ABA is essential for water stress-induced Thr-25 phosphorylation of OsMKK1.

Notably, Thr-25 was highly conserved in the OsMKK1 orthologs in maize (*Zea mays*; ZmMKK1), Arabidopsis (*Arabidopsis thaliana*; AtMKK1), soybean (*Glycine max*; GmMKK1) and *Medicago truncatula* (MtMKK1), suggesting that the phosphorylation sites of MKKs by CCaMKs are conserved in plants (Figure 4D).

### Phosphorylation of OsMKK1 at Thr-25 Enhances the Activities of OsMKK1 and OsMPK1

To determine if Thr-25 phosphorylation plays a role in OsMKK1 activation, we mutated Thr-25 of OsMKK1 either to Ala to create a non-phosphorylatable mutant (OsMKK1^T25A^) or to Asp to create a phosphomimetic mutant (OsMKK1^T25D^). In vitro kinase assays showed that the activity of OsMKK1 in the mutant OsMKK1^T25A^ did not change compared to that of OsMKK1, but the activity of OsMKK1 in the mutant OsMKK1^T25D^ had a 2.6-fold increase (Figures 4E and 4F). However, there was no difference in the autophosphorylation activity of OsMKK1 in OsMKK1, OsMKK1^T25A^ and OsMKK1^T25D^ (Figures 4E and 4F). Then, we further investigated whether Thr-25 phosphorylation regulates OsMKK1 activity in rice plants. *OsMKK1*^*T25A*^, *OsMKK1*^*T25D*^, and *OsMKK1* were over-expressed in rice under the control of the *35S* promoter. Two lines of each of *OsMKK1*^*T25A*^ (A1 and A2), *OsMKK1*^*T25D*^ (D1 and D2) and *OsMKK1*-OE (OE1 and OE2) were selected and used for further experiments, based on their increased levels of *OsMKK1* transcript and OsMKK1 protein (Supplemental Figures 15A and15B). Under the nontreated conditions, the activity of OsMKK1 in *OsMKK1*-OE1 and *OsMKK1*^*T25A*^-A1 plants was 1.6- and 1.65-fold, respectively, higher than that in WT plants, while the activity of OsMKK1 in *OsMKK1*^*T25D*^-D1 plants was 2.4-fold higher than that in WT plants (Figures 4G and 4H). After ABA treatment, OsMKK1 activity in WT, *OsMKK1*-OE1, and *OsMKK1*^*T25D*^-D1 plants increased by 2.2-, 2.5-, and 2.8-fold, respectively, while OsMKK1 activity in *OsMKK1*^*T25A*^-A1 plants increased only by 1.95-fold (Figures 4G and 4H). These results clearly indicate that Thr-25 phosphorylation in OsMKK1 is required for the activation of OsMKK1 in ABA signaling.

OsMKK1 has been reported to interact with OsMPK1 (Singh et al., 2012). To determine whether Thr-25 phosphorylation of OsMKK1 by OsDMI3 directly regulates the activity of OsMPK1, we reconstituted these three components in vitro using recombinant OsMKK1, OsMKK1^T25A^, OsMKK1^T25D^, and OsMPK1 proteins, and OsDMI3 pulled down from extracts of ABA-treated plants. After incubation with these components, the phosphorylation level of OsMPK1 was monitored. Incubation of OsMPK1 with OsMKK1 or OsMKK1^T25A^ exhibited a low phosphorylation level of OsMPK1, but incubation of OsMPK1 with OsMKK1^T25D^ significantly increased the phosphorylation level of OsMPK1 (Figures 5A and 5B). When OsDMI3 was incubated together with OsMKK1 and OsMPK1, the phosphorylation level of OsMPK1 was obviously enhanced. However, this OsDMI3-mediated phosphorylation of OsMPK1 was blocked when OsDMI3 was incubated with OsMKK1^T25A^ and OsMPK1 (Figures 5A and 5B). These results indicate that Thr-25 phosphorylation of OsMKK1 by OsDMI3 directly regulates the phosphorylation level of OsMPK1. Then, we further investigated whether Thr-25 phosphorylation regulates OsMPK1 activity in rice plants. As shown in Figures 5C and 5D, under the nontreated conditions, *OsMKK1*-OE1 and *OsMKK1*^*T25A*^-A1 plants exhibited 1.38- and 1.4-fold increases, respectively, in the activity of OsMPK1 than that of WT plants, but *OsMKK1*^*T25D*^-D1 plants showed a 1.85-fold increase of OsMPK1 activity. After ABA treatment, OsMPK1 activity in WT, *OsMKK1*-OE1, and *OsMKK1*^*T25D*^-D1 plants increased by 1.8-, 2.1-, and 2.4-fold, respectively, but OsMPK1 activity in *OsMKK1*^*T25A*^-A1 plants increased only by 1.6-fold (Figures 5C and 5D). These results indicate that Thr-25 phosphorylation of OsMKK1 positively regulates OsMPK1 activity in rice plants in ABA signaling.

**Figure 5.**
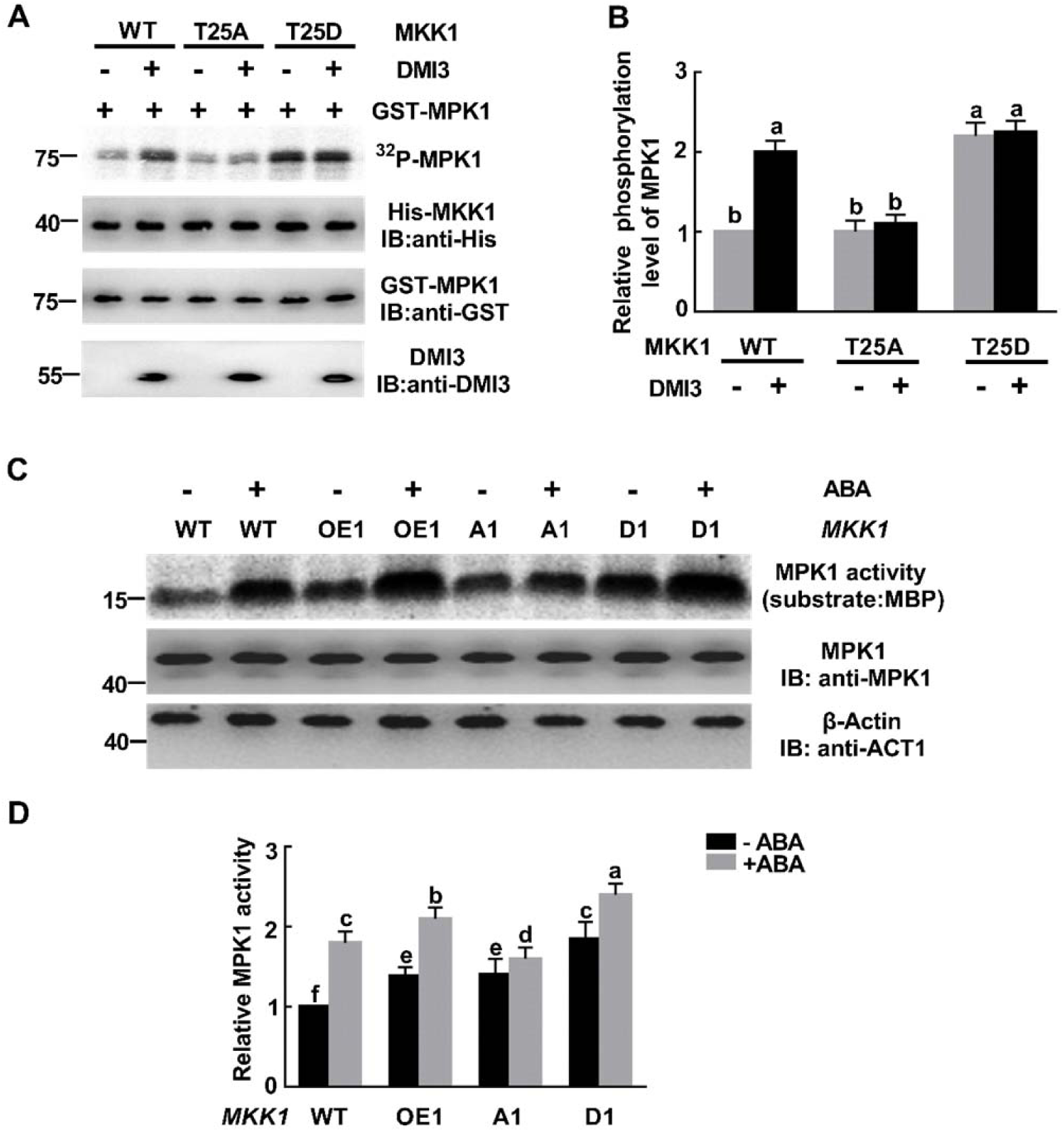
Thr-25 phosphorylation of OsMKK1 Is Required for the Activation of OsMPK1 in ABA Signaling. **(A)** In vitro reconstitution of OsDMI3-OsMKK1-OsMPK1 pathway. The recombinant OsMKK1, OsMKK1^T25A^ and OsMKK1^T25D^ were incubated with OsMPK1 in the presence or absence of OsDMI3 pulled down from extracts of ABA-treated plants, and the phosphorylation level of OsMPK1 was monitored with an in-gel kinase assay. The recombinant His-OsMKK1 was determined by IB with anti-His antibody, and the recombinant GST-OsMPK1 was determined by IB with anti-GST antibody. The amount of OsDMI3 protein was determined by IB with anti-OsDMI3 antibody. **(B)** The relative phosphorylation level of OsMPK1 in **(A)**. The phosphorylation level of OsMPK1 was quantitated by ImageJ software. The phosphorylation level of OsMPK1 incubated with OsMKK1 in the absence of OsDMI3 was set to 1. **(C)** The activation of OsMPK1 is dependent on Thr-25 phosphorylation of OsMKK1 in ABA signaling. *OsMKK1*-OE1, *OsMKK1*^*T25A*^-A1, Os*MKK1*^*T25D*^-D1, and WT plants were treated with 100 µM ABA for 2 h, and the activity of OsMPK1 was analyzed by immunoprecipitation kinase assay using MBP as substrate. OsMPK1 input was analyzed by IB with anti-OsMPK1 antibody. β-actin was used as total protein loading control. **(D)** The relative activity of OsMPK1 in **(C)**. Kinase activity was quantitated by ImageJ software. The activity of OsMPK1 in untreated WT was set to 1. All experiments were repeated at least three times with similar results. In **(B)** and **(D)**, values are means ± SEM of three independent experiments. Means denoted by the same letter did not significantly differ at P < 0.05 according to Duncan’s multiple range test. Molecular mass markers in kD were shown on the left.

Moreover, we also investigated whether Thr-25 phosphorylation of OsMKK1 by OsDMI3 could affect the interactions between OsDMI3 and OsMKK1 and between OsMKK1 and OsMPK1. Y2H assays and LCI assays showed that the interactions of OsDMI3-OsMKK1, OsDMI3-OsMKK1^T25A^, and OsDMI3-OsMKK1^T25D^ were similar (Supplemental Figures 16A and 16B) and the interactions of OsMKK1-OsMPK1, OsMKK1^T25A^-OsMPK1, and OsMKK1^T25D^-OsMPK1 were also similar (Supplemental Figures 16C and 16D), indicating that Thr-25 phosphorylation of OsMKK1 does not affect the interactions between OsDMI3 and OsMKK1 and between OsMKK1 and OsMPK1.

### The Two Modes of OsMKK1 Phosphorylation in the N-Terminus and in the Activation Loop Are Independent

In a canonical MAPK cascade, MKK is activated by MKKK through phosphorylation on two Ser/Thr residues in the conserved S/T-X5-S/T motif. In OsMKK1, the phosphorylation sites in the activation loop are Ser-215 and Thr-221 (Jagodzik et al., 2018). To test whether OsDMI3 affects the phosphorylation of OsMKK1 at Ser-215 and Thr-221 in ABA signaling, Os*MKK1-Flag* was introduced into the protoplasts of Os*DMI3*-OE1, os*dmi3*-KO1 and WT, and OsMKK1 proteins were immunoprecipitated from the transfected protoplasts with or without ABA treatment (Figure 6A). After tryptic digestion, phosphorylated peptides of Ser-215, Thr-221, and Thr-25 are quantified by parallel reaction monitoring (PRM) mass spectrometry. Experimental results showed that ABA treatment induced a significant increase in the phosphorylation of both Ser-215 and Thr-221 in the protoplasts of WT, and this ABA-induced increase was not affected in the protoplasts of both Os*DMI3*-OE1 and os*dmi3*-KO1 (Figures 6C and 6D), indicating that the phosphorylation of both Ser-215 and Thr-221 in OsMKK1 is independent of OsDMI3 in ABA signaling. At the same time, PRM analysis showed that ABA-induced Thr-25 phosphorylation of OsMKK1 is specifically dependent on OsDMI3 (Figure 6B), which is consistent with the results from the anti-phospho-Thr-25 antibody (Figure 4B).

**Figure 6.**
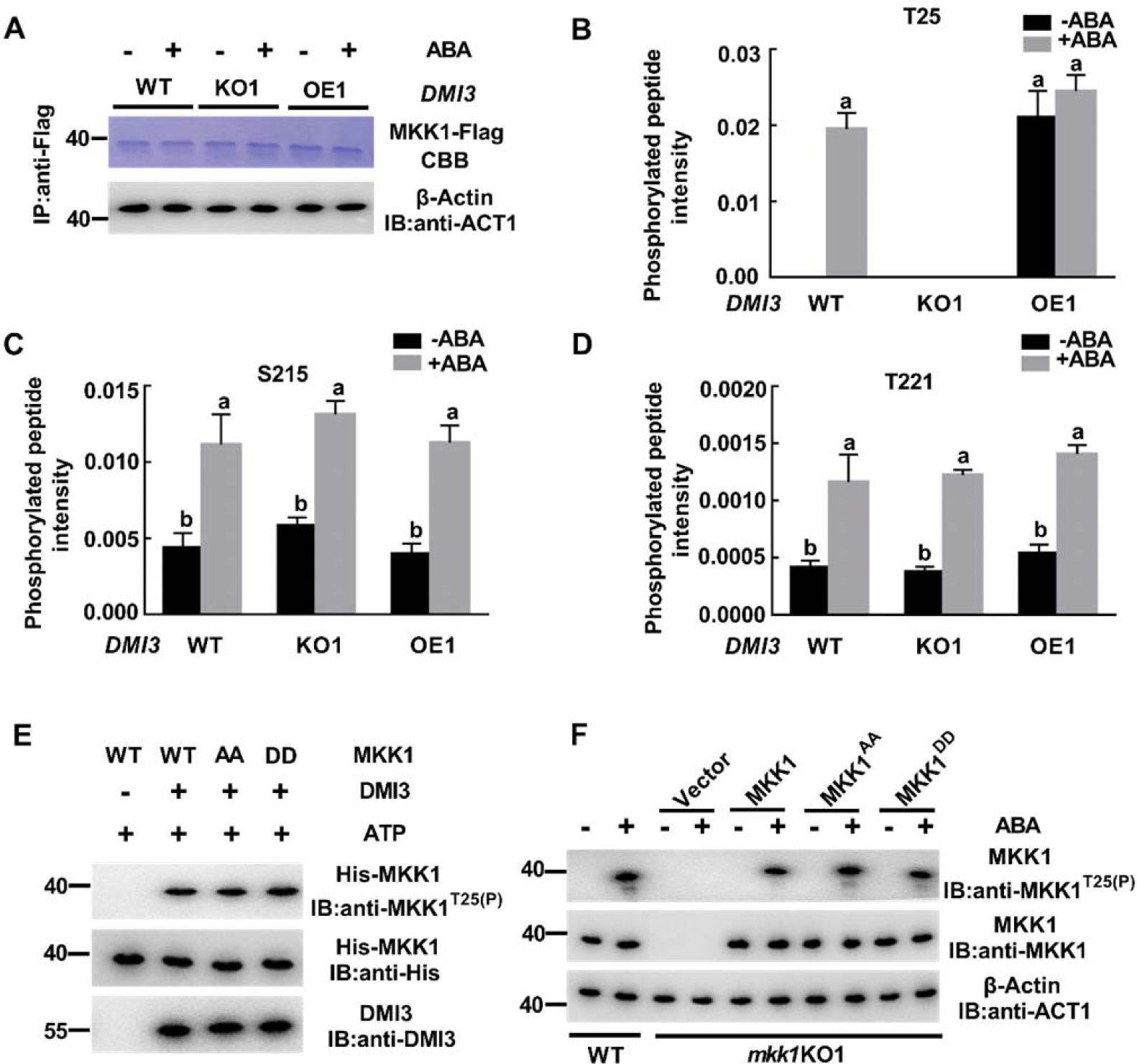
The Two Phosphorylation Pathways of OsMKK1 in the N-Terminus and in the Activation Loop Are Independent in ABA Signaling. **(A)** OsMKK1 protein immunoprecipitated from rice protoplasts. *OsMKK1-Flag* was introduced into rice protoplasts of *OsDMI3*-OE1, *osdmi3*-KO1 and WT. Transfected protoplasts were treated with 10 μM ABA for 10 min. Protein extracts were immunoprecipitated with anti-Flag antibody, followed by separation by SDS-PAGE and stained with Coomassie brilliant blue. β-actin was used as total protein loading control. **(B)** to **(D)** ABA-induced phosphorylation of Thr-25 **(B)**, Ser-215 **(C)**, and Thr-221 **(D)** in OsMKK1 in rice protoplasts. OsMKK1 protein was cut out and digested by trypsin. After tryptic digestion, the phosphorylated peptides of Thr-25, Ser-215 and Thr-221 were quantified by PRM analysis. **(E)** The phosphorylation of Thr-25 is independent of the phosphorylation of both Ser-215 and Thr-221 in vitro. Protein extracts from ABA-treated rice leaves were immunoprecipitated with anti-OsDMI3 antibody. His-OsMKK1 and mutated His-OsMKK1 (OsMKK1^S215A/T221A^ and OsMKK1^S215D/T221D^) were incubated with OsDMI3 and ATP in vitro. The phosphorylation of OsMKK1 at Thr-25 was analyzed by IB with anti-pT25 OsMKK1 antibody. The equal amount of OsDMI3 protein was shown by IB with anti-OsDMI3 antibody. His-OsMKK1 and mutated His-OsMKK1 proteins were determined by IB with anti-His antibody. **(F)** The phosphorylation of Thr-25 is not affected by the phosphorylation of both Ser-215 and Thr-221 in rice protoplasts. The indicated constructs were expressed in *osmkk1*-KO1 protoplasts, and the protoplasts were treated with 10 µM ABA for 10 min. OsMKK1 Thr-25 phosphorylation was tested by IB with anti-pT25 OsMKK1 antibody. OsMKK1 input was analyzed by IB using an anti-OsMKK1 antibody. β-actin was used as total protein loading control. All experiments were repeated at least three times with similar results. In **(B)** to **(D)**, values are means ± SEM of three independent experiments. Means denoted by the same letter did not significantly differ at P < 0.05 according to Duncan’s multiple range test. Molecular mass markers in kD were shown on the left.

Meanwhile, we also investigated whether the phosphorylation of both Ser-215 and Thr-221 affects the phosphorylation of Thr-25 in ABA signaling. Both Ser-215 and Thr-221 were mutated either to Ala (OsMKK1^S215A/T221A^) or to Asp (OsMKK1^S215D/T221D^), and *OsMKK1* and the mutant forms of *OsMKK1* were introduced into protoplasts isolated from *osmkk1*-KO1 mutant plants. The phosphorylation of OsMKK1 at Thr-25 was detected by immunoblot with the anti-phospho-Thr-25 antibody. In vitro assays showed that there was no difference in the phosphorylation of Thr-25 in OsMKK1, OsMKK1^S215A/T221A^ and OsMKK1^S215D/T221D^ in the presence of OsDMI3 and ATP (Figure 6E), indicating that the phosphorylation of both Ser-215 and Thr-221 does not affect the phosphorylation of Thr-25. Further, in vivo assays showed that under the nontreated conditions, OsMKK1 Thr-25 phosphorylation was not observed in WT protoplasts and in *osmkk1*-KO1 protoplasts transiently expressing *OsMKK1 or OsMKK1*^*S215A/T221A*^ or *OsMKK1*^*S215D/T221D*^ (Figure 6F). ABA treatment induced OsMKK1 Thr-25 phosphorylation in WT protoplasts and in *osmkk1*-KO1 protoplasts transiently expressing *OsMKK1*, and this phosphorylation was not affected in the *osmkk1*-KO1 protoplasts transiently expressing *OsMKK1*^*S215A/T221A*^ or *OsMKK1*^*S215D/T221D*^ (Figure 6F). These results indicate that the phosphorylation of Thr-25 is independent of the phosphorylation of both Ser-215 and Thr-221 in ABA signaling.

We next investigated whether the phosphorylation of both Ser-215 and Thr-221 in OsMKK1 also makes a contribution to ABA-induced activation of OsMKK1. In vitro kinase assays showed that in the OsMKK1^S215A/T221A^ mutant, the autophosphorylation and substrate phosphorylation of OsMKK1 were markedly lower than those of OsMKK1 WT, while in the OsMKK1^S215D/T221D^ mutant, the autophosphorylation and substrate phosphorylation of OsMKK1 were significantly higher than those of OsMKK1 WT (Figures 7A and 7B). Further, in vivo assays showed that in the *osmkk1*-KO1 protoplasts transiently expressing *OsMKK1*^*S215A/T221A*^, the activity of OsMKK1 was significantly lower than that of the *osmkk1*-KO1 protoplasts transiently expressing *OsMKK1* WT, but in the *osmkk1*-KO1 protoplasts transiently expressing *OsMKK1*^*S215D/T221D*^, the activity of OsMKK1 was much higher than that of the *osmkk1*-KO1 protoplasts transiently expressing *OsMKK1* WT (Figures 7C and 7D). ABA treatment induced a significant increase in the activity of OsMKK1 in the protoplasts transiently expressing *OsMKK1* WT, and the increase was further enhanced in the protoplasts transiently expressing *OsMKK1*^*S215D/T221D*^. However, this ABA-induced increase was significantly reduced in the protoplasts transiently expressing *OsMKK1*^*S215A/T221A*^ (Figures 7C and 7D). These results indicate that the phosphorylation of both Ser-215 and Thr-221 in OsMKK1 is required for ABA-induced activation of OsMKK1.

**Figure 7.**
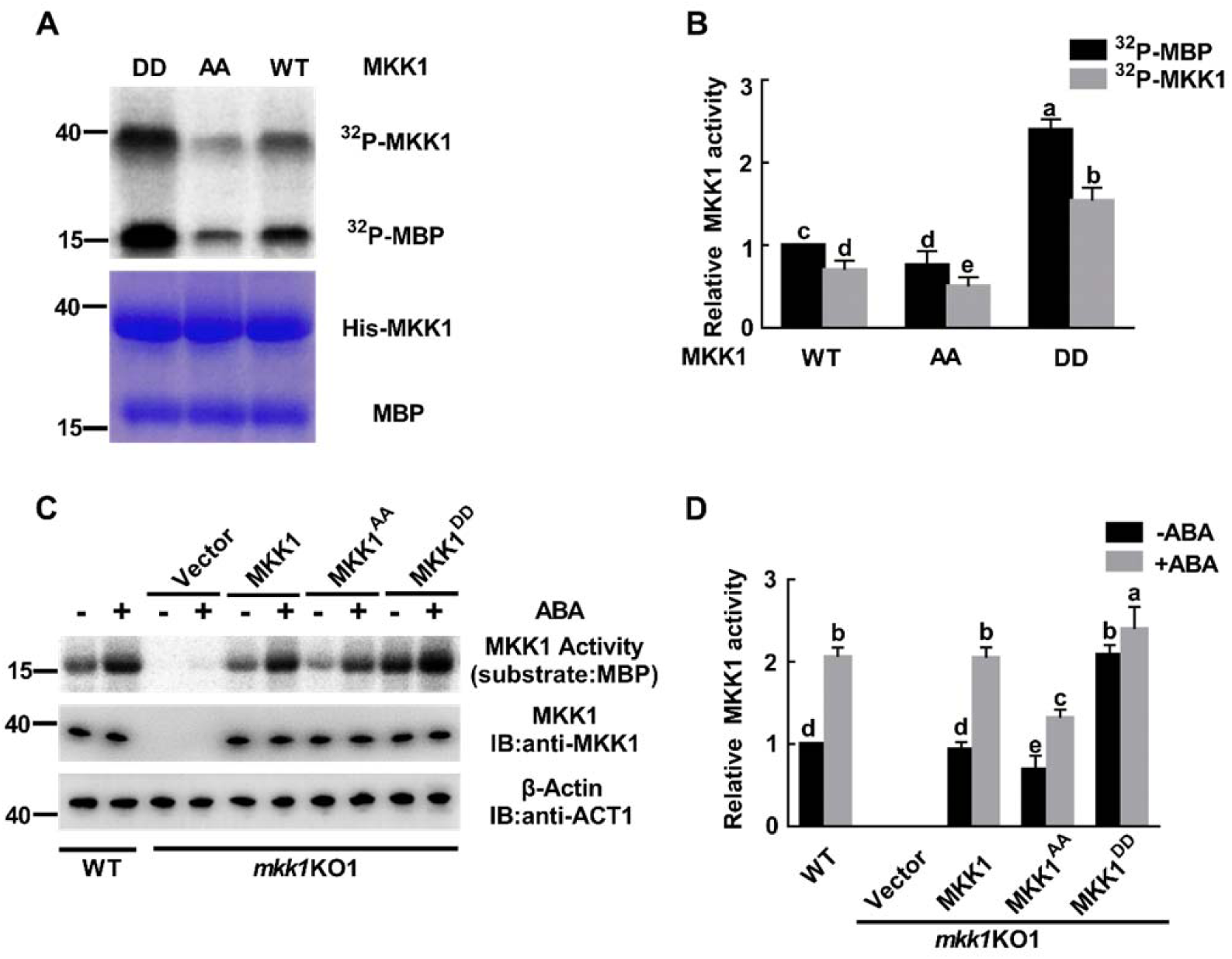
The Phosphorylation of Both Ser-215 and Thr-221 in OsMKK1 Is Required for the Activation of OsMKK1 in ABA signaling. **(A)** The activities of autophosphorylation and substrate phosphorylation of OsMKK1 in vitro. The upper gel shows substrate phosphorylation (MBP as substrate) and autophosphorylation activity of OsMKK1, OsMKK1^S215A/T221A^ and OsMKK1^S215D/T221D^, and the bottom gel shows the corresponding Coomassie staining. **(B)** The relative activities of substrate phosphorylation and autophosphorylation in **(A)**.The activities of autophosphorylation and substrate phosphorylation of OsMKK1 were quantitated by ImageJ software, and the substrate phosphorylation activity of OsMKK1 was set to 1. **(C)** OsMKK1 phosphorylation at both Ser-215 and Thr-221 is required for the activation of OsMKK1 in rice protoplasts. The indicated constructs were expressed in *osmkk1*-KO1 protoplasts, and the protoplasts were treated with 10 µM ABA for 10 min, and the activity of OsMKK1 was analyzed by immunoprecipitation kinase assay using MBP as substrate. OsMKK1 input was analyzed by IB with anti-OsMKK1 antibody. β-actin was used as the total protein loading control. **(D)** The relative activity of OsMKK1 in **(C)**. Kinase activity was quantitated by ImageJ software. The activity of OsMKK1 in untreated WT was set to 1. All experiments were repeated at least three times with similar results. In **(B)** and **(D)**, values are means ± SEM of three independent experiments. Means denoted by the same letter did not significantly differ at P < 0.05 according to Duncan’s multiple range test. Molecular mass markers in kD were shown on the left.

### OsMKK1 Positively Regulates ABA Responses and Tolerance to Both Water Stress and Oxidative Stress

Previous studies have shown that OsDMI3 is a positive regulator of ABA responses, including seed germination, root growth, antioxidant defense, and tolerance of both water stress and oxidative stress (Shi et al., 2012, 2014; Ni et al., 2019). To determine whether OsMKK1 plays a similar role to OsDMI3 in these ABA responses, we first tested the role of OsMKK1 in the regulation of ABA responses in seed germination and root growth. Under the non-treated conditions, there were no obvious differences among *OsMKK1*-OE lines (OE1 and OE2), *OsMKK1*-KO lines (KO1 and KO2), and WT plants in seed germination (Figures 8A and 8B) and primary root growth (Figures 8C and 8D). ABA treatment markedly inhibited seed germination and primary root growth in WT. The ABA sensitivity of both seed germination and primary root growth was enhanced in *OsMKK1*-OE lines, and was reduced in *osmkk1*-KO lines. These results indicate that OsMKK1, like OsDMI3, positively regulates ABA responses in seed germination and root growth.

**Figure 8.**
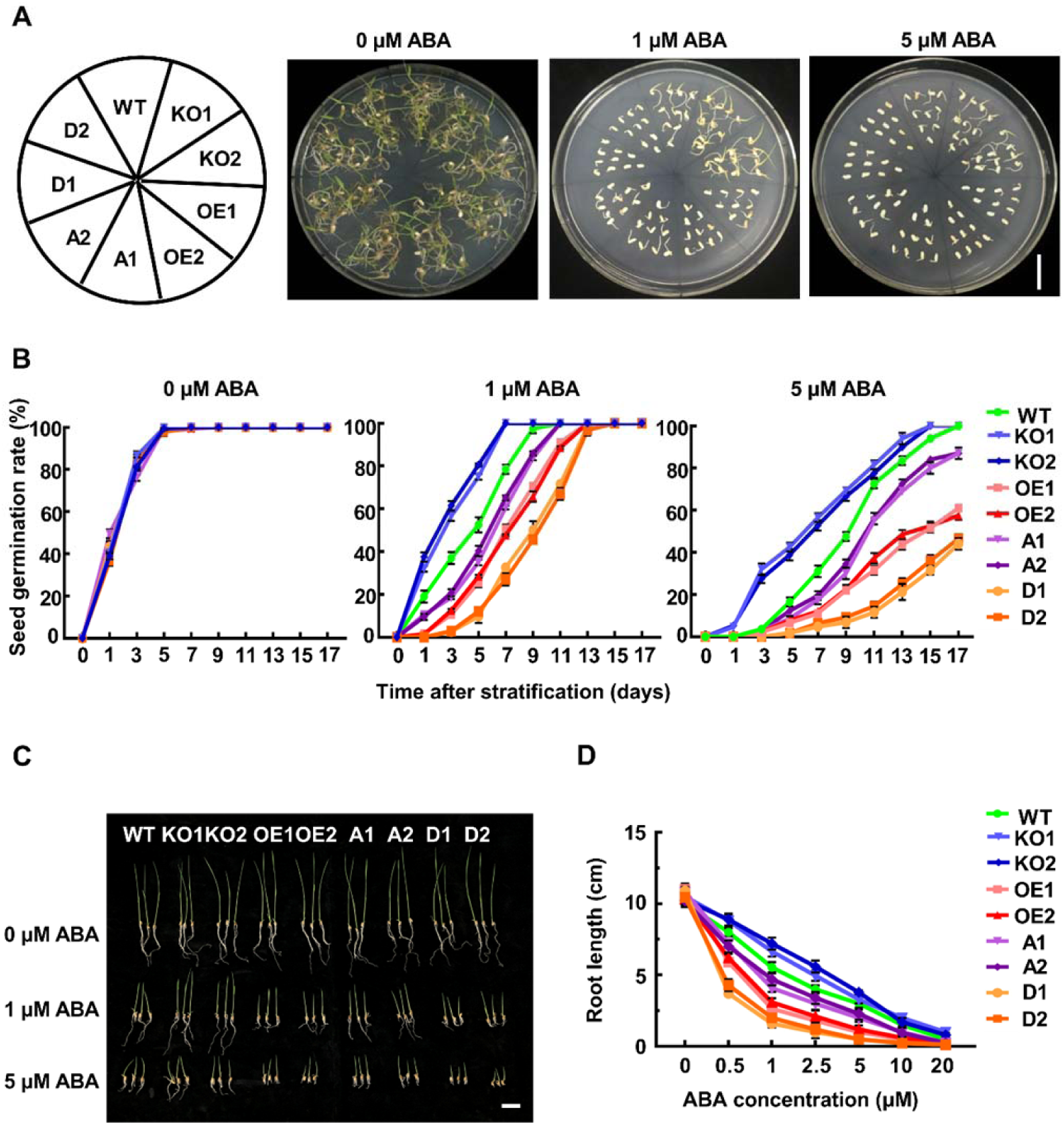
The Phosphorylation of Thr-25 Enhances ABA Sensitivity in Seed Germination and Root Growth. **(A)** Seed germination in OsMKK1-OE (OE1 and OE2), OsMKK1^T25A^ (A1 and A2), OsMKK1^T25D^ (D1 and D2), *osmkk1*-KO (KO1 and KO2) and WT. The seeds of transgenic lines and WT were germinated and grown in 1/2 MS medium supplemented with different concentrations of ABA (0, 1, 5 μM) for 9 d after stratification. Scale bar, 3 cm. **(B)** The seed germination rates of OsMKK1-OE (OE1 and OE2), OsMKK1^T25A^ (A1 and A2), OsMKK1^T25D^ (D1 and D2) and *osmkk1*-KO (KO1 and KO2) transgenic lines and WT under ABA treatments during 17 d after stratification. **(C)** Seedling growth of OsMKK1-OE (OE1 and OE2), OsMKK1^T25A^ (A1 and A2), OsMKK1^T25D^ (D1 and D2) and *osmkk1*-KO (KO1 and KO2) transgenic lines and WT under ABA treatments for 8 d. Scale bar, 4 cm. **(D)** Primary root lengths of OsMKK1-OE (OE1 and OE2), OsMKK1^T25A^ (A1 and A2), OsMKK1^T25D^ (D1 and D2), *osmkk1*-KO (KO1 and KO2) transgenic lines and WT grown in different concentrations of ABA as indicated for 10 d. Approximately 48 seeds of each transgenic line were analyzed per replicate for each concentration of ABA in **(A)** to **(D)**. In **(A)** and **(C)**, experiments were repeated at least three times with similar results. In **(B)** and values are means ± SEM of three independent experiments.

We next investigated whether OsMKK1 is involved in the tolerance of water stress and oxidative stress in rice plants. *OsMKK1*-OE, *osmkk1*-KO, and WT plants were treated with either PEG to simulate water stress or H_2_O_2_ to produce oxidative stress, and the phenotype of these plants under the stressed conditions was analyzed. Under the non-stressed conditions, there was no obvious morphological difference between the transgenic plants and the WT plants (Figures 9A and 9B). When treated with 20% PEG (Figure 9A) or with 100 mM H_2_O_2_ (Figure 9B), the *osmkk1*-KO plants exhibited more sensitive to water stress and oxidative stress compared with WT plants, and had lower survival rates after recovery by re-watering (Figures 9C and 9D). In contrast, the *OsMKK1*-OE plants showed enhanced tolerance to water stress and oxidative stress after PEG and H_2_O_2_ treatments and had higher survival rates after recovery by re-watering. Consistent with the phenotype of these transgenic plants under the stressed conditions, the *osmkk1*-KO plants had a higher oxidative damage, indicated by the content of malondialdehyde (MDA) (Figure 9E) and the percentage of electrolyte leakage (Figure 9F), than WT plants under the stressed conditions, whereas the *OsMKK1*-OE plants had a lower oxidative damage than WT plants. These results indicate that OsMKK1 positively regulates the tolerance of rice plants to water stress and oxidative stress.

**Figure 9.**
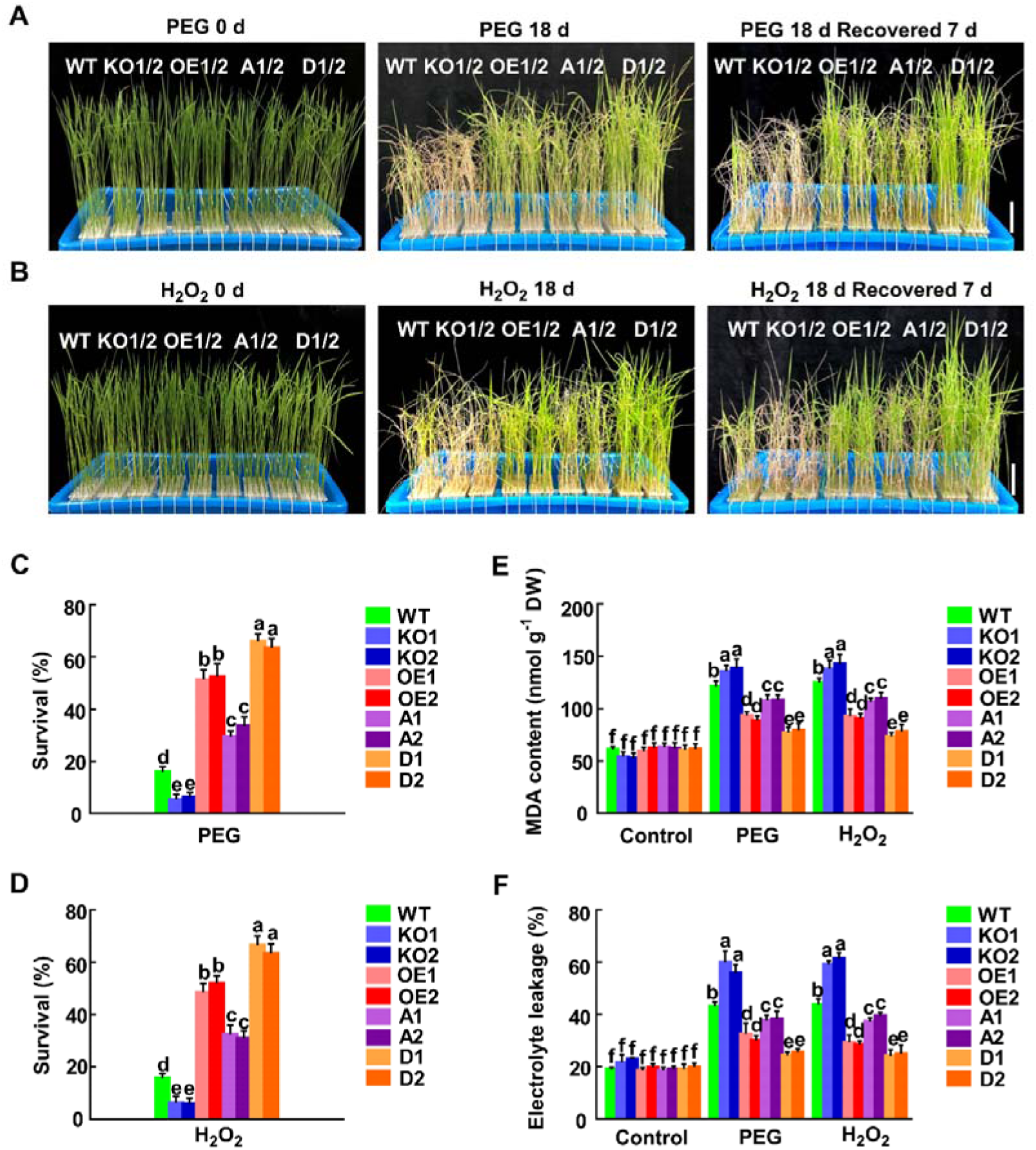
The Phosphorylation of Thr-25 Positively Regulates the Tolerance of Rice Plants to Water Stress and Oxidative Stress. **(A)** and **(B)** Phenotypes of OsMKK1-OE (OE1 and OE2), OsMKK1^T25A^ (A1 and A2), OsMKK1^T25D^ (D1 and D2), *osmkk1*-KO (KO1 and KO2) and WT plants exposed to water stress **(A)** or oxidative stress **(B)**. Ten-day-old rice seedlings were treated with 20% PEG 4000 **(A)** or 100 mM H_2_O_2_ **(B)** for 18 d, and then recovered by re-watering for 7 d. Approximately 40 rice seedlings of each transgenic line were used per replicate. Scale bars, 4.5 cm. **(C)** and **(D)** The survival rate (%) of the rice plants exposed to water stress **(C)** or oxidative stress **(D)** after recovery by re-watering for 7 d shown in **(A)** and **(B)**. **(E)** and **(F)** The content of MDA **(E)** and the percent leakage of electrolyte **(F)** in the leaves of OsMKK1-OE (OE1 and OE2), OsMKK1^T25A^ (A1 and A2), OsMKK1^T25D^ (D1 and D2), *osmkk1*-KO (KO1 and KO2) and WT plants exposed to water stress or oxidative stress. Ten-day-old seedlings were treated with 20% PEG 4000 or 100 mM H_2_O_2_ for 2 d, and then leaves were sampled for the determination of MDA content and electrolyte leakage (%). Approximately 40 seedlings of each transgenic line were used per replicate. In **(A)** and **(B)**, experiments were repeated at least three times with similar results. In **(C)** to **(F)**, values are means ± SEM of three independent experiments. Means denoted by the same letter did not differ significantly at P < 0.05 according to Duncan’s multiple range test.

### Phosphorylation of OsMKK1 by OsDMI3 Enhances ABA Sensitivity and Tolerance to Both Water Stress and Oxidative Stress

To test whether Thr-25 phosphorylation of OsMKK1 contributes to the function of OsMKK1 in ABA signaling, *OsMKK1*-OE (OE1 and OE2), *OsMKK1*^*T25A*^ (A1 and A2), *OsMKK1*^*T25D*^ (D1 and D2), and WT were treated with ABA, PEG and H_2_O_2_, respectively, and the phenotype of these plants under the stressed conditions was analyzed. Under the non-treated conditions, there were no obvious differences between the transgenic lines and the WT in seed germination (Figures 8A and 8B), primary root growth (Figures 8C and 8D), and the growth of seedlings (Figures 9A and 9B). After ABA treatment, there were significant differences among *OsMKK1*-OE, *OsMKK1*^*T25A*^, and *OsMKK1*^*T25D*^ lines in seed germination (Figures 8A and 8B) and primary root growth (Figures 8C and 8D), in which *OsMKK1*^*T25D*^ lines exhibited a significantly enhanced sensitivity to ABA compared with *OsMKK1*-OE lines, but *OsMKK1*^*T25A*^ lines displayed a markedly reduced sensitivity to ABA, indicating that Thr-25 phosphorylation makes an important contribution to the function of OsMKK1 in these ABA responses. Further, *OsMKK1*^*T25D*^ plants exhibited enhanced tolerance to water stress (Figure 9A) and oxidative stress (Figure 9B) compared with *OsMKK1*-OE plants, with a higher survival rate after recovery by re-watering (Figures 9C and 9D) and a lower oxidative damage (Figures 9E and 9F). By contrast, *OsMKK1*^*T25A*^ plants showed reduced tolerance to water stress and oxidative stress, with a lower survival rate after recovery by re-watering and a higher oxidative damage. These results indicate that Thr-25 phosphorylation plays an important role in OsMKK1-regulated tolerance of rice plants to water stress and oxidative stress.

To further confirm that it is indeed the OsDMI3-OsMKK1 pathway that regulates ABA response and tolerance to water stress and oxidative stress in rice plants, the mutants *osdmi3-*KO/*OsMKK1*-OE, *osdmi3-*KO/*OsMKK1*^*T25A*^, *osdmi3-*KO/*OsMKK1*^*T25D*^ (Supplemental Figures 15C and 15D), *osmkk1*-KO/*OsDMI3*-OE (Supplemental Figures 15E and 15F), and *osmkk1*-KO/*osdmi3-*KO (Supplemental Figure 15G) were generated, and the phenotypes of these transgenic lines under ABA treatment and stress conditions were analyzed. Compared with the *OsDMI3*-OE line, the *osmkk1*-KO/*OsDMI3*-OE line exhibited a dramatic reduction in the sensitivity of seed germination to ABA (Figures 10A and 10B) and in the tolerance to water stress (Figure 10C) and oxidative stress (Figure 10D), with a lower survival rate (Figures 10E and 10F), a higher oxidative damage (Supplemental Figures 17A and 17B), and a faster water loss (Supplemental Figure 17C). The sensitivity of the *osmkk1*-KO/*OsDMI3*-OE line to ABA and the tolerance to water stress and oxidative stress were close to, but still significantly higher than those of the *osmkk1* line. These results suggest that OsDMI3 mediates these responses partly through the action of OsMKK1. On the other hand, the sensitivity of the *osdmi3*-KO/*OsMKK1*-OE line to ABA and the tolerance to water stress and oxidative stress are much lower than those of the *OsMKK1*-OE line but significantly higher than those of the *osdmi3*-KO line, suggesting that the role of OsMKK1 in these responses is partly dependent on OsDMI3. Further, the *osdmi3*-KO/*OsMKK1*^*T25A*^ line displayed a similar phenotype to the *osdmi3*-KO/*OsMKK1*-OE line in its sensitivity to ABA and tolerance to water and oxidative stress, but the *osdmi3-*KO/*OsMKK1*^*T25D*^ line exhibited a greatly enhanced sensitivity to ABA and a greatly enhanced tolerance to water stress and oxidative stress, indicating that OsDMI3-mediated Thr-25 phosphorylation of OsMKK1 plays an important role in the ABA responses. Moreover, compared with the *osdmi3*-KO line and the *osmkk1*-KO line, the *osmkk1*-KO/*osdmi3*-KO double mutant showed less sensitivity to ABA and lower tolerance to water stress and oxidative stress, suggesting that, in addition to the OsDMI3-OsMKK1 pathway, there are other targets of OsDMI3 and other activation pathway of OsMKK1 involved in the ABA responses. Taken together, these results support the OsDMI3-OsMKK1 pathway is involved in the regulation of ABA responses.

**Figure 10.**
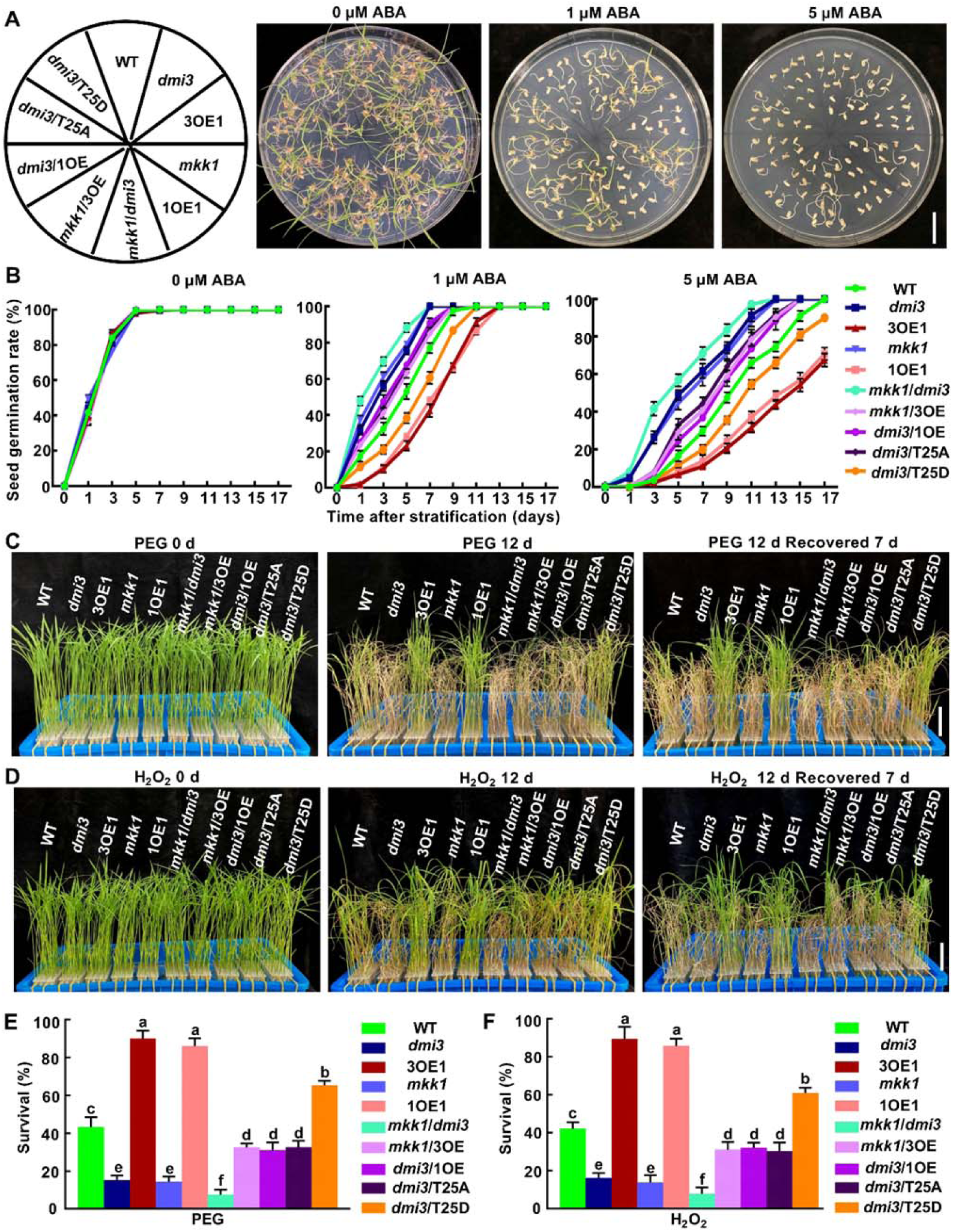
The OsDMI3-OsMKK1 Pathway Regulates ABA Response and Tolerance to Water Stress and Oxidative Stress. **(A)** Seed germination in *osdmi3-*KO1 (*dmi3*), *OsDMI3*-OE1 (3OE1), *osmkk1*-KO1 (*mkk1*), *OsMKK1*-OE1 (1OE1), *osmkk1*-KO/*osdmi3-*KO *(mkk1*/*dmi3), osmkk1*-KO/*OsDMI3*-OE (*mkk1*/3OE), *osdmi3-*KO/*OsMKK1*-OE (*dmi3*/1OE), *osdmi3-*KO/*OsMKK1*^*T25A*^ (*dmi3*/T25A), *osdmi3-*KO/*OsMKK1*^*T25D*^ (*dmi3*/T25D), and WT. The seeds of transgenic lines and WT were germinated and grown in 1/2 MS medium supplemented with different concentrations of ABA (0, 1, 5 μM) for 12 d after stratification. Approximately 40 rice seedlings of each transgenic line were used per replicate for each concentration of ABA. Scale bar, 3 cm. **(C)** The seed germination rates of the transgenic lines shown in **(A)** and WT under ABA treatments during 17 d after stratification. **(C)** and **(D)** Phenotypes of the transgenic plants shown in **(A)** and WT plants exposed to water stress **(C)** or oxidative stress **(D)**. Ten-day-old rice seedlings were treated with 20% PEG 4000 **(C)** or 100 mM H_2_O_2_ **(D)** for 12 d, and then recovered by re-watering for 7 d. Approximately 40 rice seedlings of each transgenic line were used per replicate. Scale bars, 4.5 cm. **(E)** and **(F)** The survival rate (%) of the rice plants exposed to water stress **(E)** or oxidative stress **(F)** after recovery by re-watering for 7 d shown in **(C)** and **(D)**. In **(A), (C)**, and **(D)**, experiments were repeated at least three times with similar results. In **(B), (E)**, and **(F)**, values are means ± SEM of three independent experiments. In **(E)** and **(F)**, means denoted by the same letter did not differ significantly at P < 0.05 according to Duncan’s multiple range test.

## DISCUSSION

CCaMK (DMI3) has been shown to be an important positive regulator of ABA and abiotic stress signaling in plants (Shi et al., 2012, 2014; Zhu et al., 2016b; Ni et al., 2019). A double-negative regulatory pathway for the activation of the rice CCaMK OsDMI3 in ABA signaling has been established, whereby the type 2C protein phosphatase OsPP45 inactivates OsDMI3 by direct dephosphorylation in the absence of ABA, but in the presence of ABA, ABA-induced H_2_O_2_ inhibits OsPP45 activity, resulting in OsDMI3 activation (Ni et al., 2019). Once activated, CCaMK can phosphorylate its protein targets. In maize, the NAC (NAM, ATAF1/2, and CUC2) transcription factor ZmNAC84 has been identified to be a direct target of ZmCCaMK (Zhu et al., 2016b). ZmNAC84 interacted with and was phosphorylated by ZmCCaMK, and the phosphorylation at Ser-113 of ZmNAC84 by ZmCCaMK is essential for ABA-induced antioxidant defense. Moreover, it was also shown that OsDMI3 and OsMPK1 are in the same signaling pathway in ABA signaling, in which OsDMI3 functions upstream of OsMPK1 to regulate the activities of antioxidant enzymes and the production of H_2_O_2_ (Shi et al., 2014). Previous studies have demonstrated that OsMPK1 and its orthologs in other plants, such as AtMPK6 in Arabidopsis and SIPK (salicylic acid-induced protein kinase) in tobacco (*Nicotiana tabacum*), are the key components involved in ABA and abiotic stress signaling in plants (Colcombet and Hirt, 2008; Xing et al., 2008; Šamajová et al., 2013; Singh and Jwa, 2013; Danquah et al., 2014; de Zelicourt et al., 2016; Dóczi and Bögre, 2018; Jagodzik et al., 2018). However, it is unknown whether OsMPK1 is a direct target of OsDMI3 in ABA signaling and, If not, how OsDMI3 induces the activation of OsMPK1 in ABA signaling.

In this study, we identified that OsDMI3 does not interact directly with OsMPK1 (Supplemental Figure 1), but interacts with the OsMPK1 activators OsMKK1 and OsMKK6 (Figure 1; Supplemental >Figure 2). We found that OsDMI3 is required for ABA-induced phosphorylation and activation of OsMKK1 but not OsMKK6 (Figures 2 and 3). We further identified that OsDMI3 directly phosphorylates OsMKK1 at Thr-25 (Figure 4A; Supplemental Figure 12), and this Thr-25 phosphorylation is OsDMI3-specific in ABA signaling (Figure 4B) and ABA-specific under water stress (Figure 4C). Our genetic evidence indicated that OsMKK1 is a positive regulator of ABA responses, including seed germination, root growth, and tolerance of both water stress and oxidative stress, and the phosphorylation of OsMKK1 by OsDMI3 makes an important contribution to the function of OsMKK1 in the ABA responses (Figures 8 to 10).

In MAPK signaling, the interactions between MAPKs and their activators, substrates, and inactivators are commonly achieved through specific docking interactions (Bardwell, 2006; Bigeard and Hirt, 2018; Krysan and Colcombet, 2018). The docking interactions mediated by a common docking domain (CD domain) in the C-terminal region of MAPKs and a docking domain (D-domain) in the N-terminal tail of MKKs have been extensively characterized in animals and plants. D-domains can also be found in MAPK regulatory proteins and substrates. However, the protein domains involved in MKKK-MKK interactions have been less well characterized. In mammalian cells, it was found that the C termini of MEK1, MKK3, MKK4, MKK6, and MKK7 contain a conserved docking site, termed DVD (domain for versatile docking), which binds to the kinase domain of their specific upstream MKKKs, and such docking interactions are required for the activation of these MKKs (Takekawa et al., 2005). In Arabidopsis, it was shown that the C-terminal region of MKKK20 is required for the interaction with MKK3 (Bai and Matton, 2018). Surprisingly, only the full-length MKK3 interacted with MKKK20, the N-terminus, kinase domain and the C-terminus of MKK3 did not interact with MKKK20. In this study, we found that among the six rice MKKs we tested (OsMKK1, OsMKK6, OsMKK3, OsMKK4, OsMKK5, and OsMKK10-2), only the group A MKKs, OsMKK1 and OsMKK6, interact with OsDMI3 (Figure 1; Supplemental Figure 2), and the interactions are mediated by the EF hands domain in the C-terminus of OsDMI3 and the N-terminus domain of OsMKK1 or OsMKK6 (Supplemental Figures 3 to 6). The interaction domains between OsDMI3 and OsMKKs are distinct from those between MKKKs and MKKs in animal systems (Takekawa et al., 2005), suggesting that the docking domain in MKKs for MKKKs is different from that in MKKs for their regulatory proteins.

Interestingly, although both OsMKK1 and OsMKK6 can be phosphorylated by OsDMI3 in vitro, only ABA-induced phosphorylation and activation of OsMKK1 are OsDMI3-dependent, while ABA-induced phosphorylation and activation of OsMKK6 are OsDMI3-independent (Figures 2 and 3). These results indicate that OsMKK1 is and OsMKK6 is not a physiological substrate of OsDMI3. This opens up the interesting question of why OsDMI3 cannot phosphorylate OsMKK6 in ABA signaling. In yeast and animal systems, multiple mechanisms have been shown to be involved in the specificity of protein phosphorylation, including (1) the structures of the kinase and the substrate, (2) local and distal interactions between the kinase and the substrate, (3) the formation of complexes with scaffolding and adaptor proteins that spatially regulate the kinase, and (4) systems-level effects such as competition, multisite phosphorylation and kinetic proofreading (Ubersax and Ferrell, 2007; Miller and Turk, 2018). Plants may employ similar mechanisms to maintain the specificity of protein phosphorylation (Pitzschke, 2015; Bigeard and Hirt, 2018; Dóczi and Bögre, 2018; Krysan and Colcombet, 2018). Because no crystal structures of OsDMI3, OsMKK1 and OsMKK6 have yet been elucidated, we do not know whether there exists a structural constraint between OsDMI3 and OsMKK6, which blocks OsDMI3 to phosphorylate OsMKK6 in ABA signaling. In addition, there might exist a systems-level competition between OsMKK1 and OsMKK6, which may explain why OsDMI3 fails to phosphorylate OsMKK6 in ABA signaling.

A canonical MAPK cascade is composed of three sequentially activating kinases: a MKKK phosphorylates and activates a MKK on two Ser and/or Thr residues in a conserved (S/T)-X5-(S/T) motif in plants and (S/T)-X3-(S/T) in yeast and animals (Ren et al., 2002; Hamel et al., 2012), which then phosphorylates a MAPK by dual phosphorylation of the conserved TXY motif located in its activation loop (Hamel et al., 2012; Danquah et al., 2014; Jagodzik et al., 2018). This three-component MAPK cascade is highly conserved in eukaryotes (Rodriguez et al., 2010; Hamel et al., 2012; >Bigeard and Hirt, 2017; Jagodzik et al., 2018). However, some non-canonical MAPK pathways have also been reported in both animals and plants. The mammalian MAPK p38α has been shown to be activated by direct interaction with the protein TAB1 [transforming growth factor-β-activated protein kinase 1 (TAK1)-binding protein 1] (Ge et al., 2002) or through Tyr-323 phosphorylation by T cell receptor-proximal tyrosine kinases (Salvador et al., 2005), and both pathways induce the cis-autophosphorylation of its activation loop. In plants, several different regulatory mechanisms for the MKK-independent activation of MAPKs have been described. In a salt stress response, Arabidopsis AtMPK6 can be activated by the binding of phosphatidic acid (Yu et al., 2010). Under mechanical wounding, the full activation of AtMPK8 requires direct binding of CaMs in a Ca^2+^-dependent manner in addition to MKK3 phosphorylation (Takahashi et al., 2011). In an immune response, the rice CDPK OsCPK18 is able to activate OsMPK5 by phosphorylating two conserved Thr residues (Thr-14 and Thr-32) in the N-terminal region of OsMPK5 without affecting the phosphorylation of its TXY motif (Xie et al., 2014). Moreover, a recent study showed that the transmembrane kinases (TMKs) TMK1 and TMK4 directly and specifically interact with and phosphorylate MKK4/5, which is required for auxin to activate MKK4/5-MPK3/6 signaling (Huang et al., 2019). However, the phosphorylation sites of MKK4/5 by these TMKs were not identified. Therefore, it is not clear whether or not these TMKs are MKKK-like kinases. In this study, we show an OsDMI3-dependent activation pathway of OsMKK1 in rice, which is distinct from these reported examples of unconventional MAPK activation in plants and animals. We found that OsDMI3 directly phosphorylates Thr-25 in the N-terminus of OsMKK1, but not the two Ser/Thr residues (Ser-215/Thr-221) in the conserved S/T-X5-S/T motif (Figures 4A to 4C), and this phosphorylation does not affect the autophosphorylation of OsMKK1 (Figures 4E and 4F). The activation of OsMKK1 (Figure 4) and OsMPK1 (Figure 5) is dependent on Thr-25 phosphorylation of OsMKK1 by OsDMI3 in ABA signaling. Our data indicate that OsDMI3-mediated phosphorylation of OsMKK1 is an important regulatory mechanism for OsMPK1 activation in ABA signaling (Figure 11).

**Figure 9.**
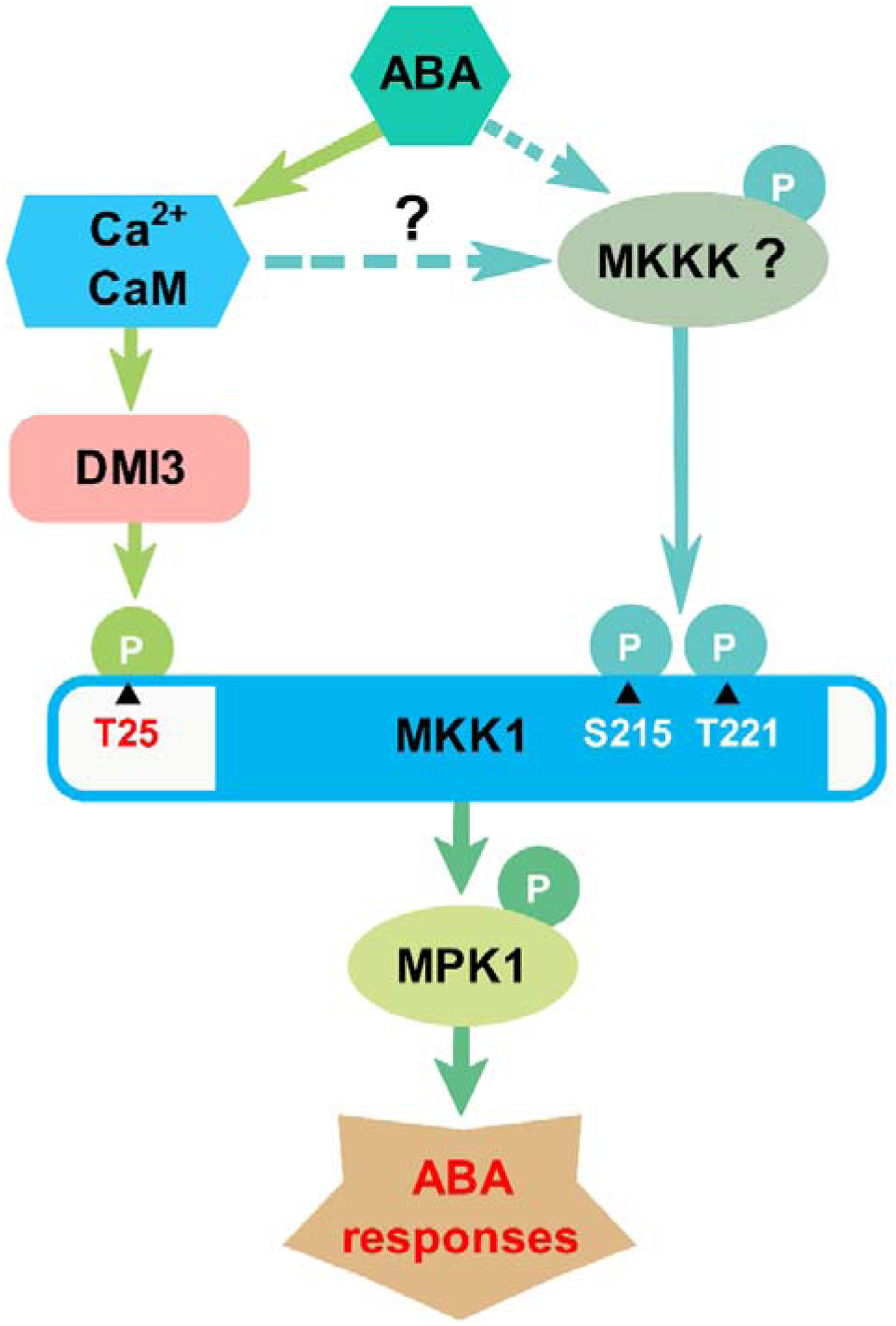
A Working Model for OsDMI3-Mediated Activation of MAPK Cascade in ABA Signaling. ABA induces the activation of OsDMI3, and the activated OsDMI3 directly phosphorylates OsMKK1 at Thr-25 to enhance its activity, thus resulting in the activation of OsMPK1. On the other hand, ABA also induces the phosphorylation of both Ser-215 and Thr-221 in OsMKK1, which are the phosphorylation sites in the activation loop of OsMKK1 by OsMKKK, showing that an OsMKKK-dependent pathway is involved in the regulation of OsMKK1 activity in ABA signaling. The two modes of OsMKK1 phosphorylation are independent and the OsDMI3-mediated phosphorylation of OsMKK1 makes a major contribution to the activation of OsMKK1 and the function of OsMKK1 in ABA signaling. However, the OsMKKK for the ABA-induced activation of the OsMKK1-OsMPK1 cascade is unknown. Moreover, it is also unclear whether a Ca^2+^/CaM-dependent pathway functions upstream of the canonical MAPK cascade in ABA signaling.

Interestingly, the N-terminal Thr-25 residue of OsMKK1 is conserved in the monocots and dicots, such as maize, Arabidopsis, soybean, and *Medicago truncatula* (Figure 4D). In legumes, CCaMKs have been shown to play key roles in mediating symbiotic relationships with rhizobial and arbuscular mycorrhizae (Singh and Parniske, 2012; Poovaiah et al., 2013). A recent study by quantitative phosphoproteomic analyses suggested that MAPK-mediated phosphorylation signaling may be involved in the rhizobia–legume symbiosis (Zhang et al., 2019). It is possible that the CCaMK-MKK-MAPK pathway might also be involved in the establishment of symbiotic relationships.

However, in this study, our results also showed that ABA-induced phosphorylation and activation of OsMKK1 were not fully inhibited in the *osdmi3* mutant (Figures 2E, 2F, 3C, and 3D). Our genetic results showed that the role of OsMKK1 in ABA responses is only partly dependent on OsDMI3 (Figure 10). These results indicate that an OsDMI3-independent pathway is also involved in the activation of OsMKK1 in ABA signaling. Further, we found that ABA treatment not only induces an increase in the phosphorylation of OsMKK1 Thr-25, but also induces an increase in the phosphorylation of both Ser-215 and Thr-22 (Figures 6B to 6D), which are the phosphorylation sites in the activation loop of OsMKK1 by OsMKKK (Jagodzik et al., 2018), and the phosphorylation of both Ser-215 and Thr-221 in OsMKK1 is also essential for the activation of OsMKK1 in ABA signaling (Figure 7), suggesting that OsMKKK is involved in the activation of OsMKK1 in ABA signaling. However, this ABA-induced increase in the phosphorylation of both Ser-215 and Thr-221 is not affected by OsDMI3. On the other hand, the phosphorylation of Thr-25 is also independent of the phosphorylation of both Ser-215 and Thr-221 in ABA signaling (Figures 6E and 6F). Taken together, our results suggest that the phosphorylation of OsMKK1 by both OsDMI3 and OsMKKK is independent, and the full activation of OsMKK1 requires both OsDMI3 and OsMKKK in ABA signaling (Figure 11).

In Arabidopsis, several complete MAPK cascades, such as MAPKKK17/18-MKK3-MPK1/2/7/14 (Danquah et al., 2015; Matsuoka et al., 2015) and MAPKKK20-MKK5-MPK6 (Li et al., 2017), have been shown to be involved in ABA signaling. However, the exact OsMKKK for the ABA-induced activation of the OsMKK1-OsMPK1 cascade is unknown. Moreover, in this study, Ca^2+^ was shown to be required for ABA-induced activation of OsMKK1, but Ca^2+^-induced increase in OsMKK1 activity was not completely blocked in *osdmi3*-KO1 plants (Supplemental Figure 9), indicating that the other Ca^2+^-dependent pathway is also involved in ABA-induced activation of OsMKK1. Accumulating data have demonstrated that the receptor-like protein kinases/receptor-like proteins are key regulators of MAPK cascades (Xu and Zhang, 2015; Zhang et al., 2018). CRLK1, a Ca^2+^/CaM-regulated receptor-like kinase, has been reported to play a critical role in plant response to cold stress (Yang et al., 2010a; Zhao et al., 2017). CRLK1 physically interacts with and phosphorylates MEKK1 (Yang et al., 2010b; Furuya et al., 2013). CRLK1 and CRLK2 positively regulate cold response, possibly by activating the MEKK1-MKK2-MPK4 pathway and by suppressing the cold activation of the MKK4/5-MPK3/6 pathway (Zhao et al., 2017). However, whether such a Ca^2+^/CaM-regulated receptor-like kinase is involved in the ABA-activated OsMKKK-OsMKK1-OsMPK1 cascade remains to be determined (Figure 11).

## METHODS

### Plant Materials and Constructs

Rice (*Oryza sativa*) plants used in this study include Nipponbare (WT), *osdmi3*, and *OsDMI3* (Ni et al., 2019). Rice seedlings were grown under the conditions as described previously (Zhang et al., 2014). When the second leaves were fully expanded, they were collected and used for various investigations.

All transgenic lines, including the overexpressing (OE) lines of *OsMKK1, OsMKK6*, and *OsABA2*, the knockout lines (KO) of *osmkk1, osmkk6*, and *osaba2*, the *osdmi3*-KO mutant plants overexpressing *OsMKK1* (*osdmi3*/*OsMKK1*), *OsMKK1*^*T25A*^ (*osdmi3*/*OsMKK1*^*T25A*^), and *OsMKK1*^*T25D*^ (*osdmi3*/*OsMKK1*^*T25D*^), the *osmkk1*-KO mutant plants overexpressing *OsDMI3* (*osmkk1*/*OsDMI3*), and the double mutant *osmkk1*/*osdmi3*, were generated by Biogel Company (Hangzhou, China). To generate *OsMKK1*, Os*MKK6, OsMKK1*^*T25A*^, *OsMKK1*^*T25D*^, *OsABA2*, and *OsDMI3* constructs, *OsMKK1, OsMKK6* and the mutated *OsMKK1* coding sequences were inserted into the pCAMBIA 1304 vector by *Nco*I and *Spe*I sites, the coding sequence of *OsDMI3* were inserted into the pCAMBIA 1304 vector by *Kpn*I and *Spe*I sites, and the coding sequence of *OsABA2* were inserted into the pCAMBIA 1304 vector by *Nco*I and *Kpn*I sites. The constructs were introduced into *Agrobacterium* strain EHA105 and transformed into rice (*O. sativa* sub. *japonica* cv Nipponbare) under the control of CaMV 35S promoter. The transgenic plants were selected on 50 μg mL^-1^ hygromycin. The homozygous T3 seeds of transgenic plants were used for further analysis.

To generates the *osmkk1, osmkk6, osaba2*, and *osmkk1*/*osdmi3* mutant plants, the CRISPR/Cas9 system was used. The sgRNAs of *OsMKK1, OsMKK6, OsABA2, and OsDMI3* were shown in Supplemental Figures 7, 14 and 15G. The single sgRNA was created in the BGK03 vector containing Cas9. The constructs were introduced into rice (*O. sativa* sub. *japonica* cv Nipponbare) by *Agrobacterium*-mediated transformation method. Positive transgenic individuals were identified by sequencing analyses (for primers, Supplemental Table 3).

### Y2H Assay

Y2H analysis was performed using the Matchmaker Gold Yeast Two-Hybrid System (Clontech) according to the manufacturer’s protocol. Full-length *OsMKK1/OsMKK6* were separately cloned into pGADT7 vector at the *Sma*I-*BamH*I sites and the *EcoR*I-*Sma*I sites, and the constructs were transformed into the yeast strain Y187. The transformed Y187 yeast strain was mated with the Y2HGold yeast strain containing pGBKT7-OsDMI3 (Ni et al., 2019). The mating yeast cultures were spread on plates (SD/-Trp/-Leu/-His/-Ade) containing X-α-gal (40 μg mL^-1^) and AbA (0.125 μg mL^-1^). To determine the interaction intensity, dilutions of yeast cultures (10^-0^, 10^-1^, and 10^-2^) were spotted onto selection medium for blue color development. Photographs were taken after 5-7 days incubation at 30°C.

### GST Pull-Down Assay

OsDMI3-GST fusion protein was produced in *Escherichia coli Rosetta* (*DE3*; Novagen), and tested for interaction as described previously (Ni et al., 2019). Full-length coding sequences of *OsMKK1*/*OsMKK6* were cloned into the pET30a vector (Novagen) at the *Kpn*I-*BamH*I sites to generate His tag fusion proteins. Expression of His-OsMKK1/OsMKK6 in *Escherichia coli Rosetta* (*DE3*) cells was induced with 0.5 mM isopropyl-β-D-thiogalactoside at 24°C for 6 h. Fusion proteins were purified using MagnetHis™ protein purification system (Promega) according to the manufacturer’s protocol.

For pull-down assay, GST or GST-OsDMI3 immobilized on the Magnet GST Particles (Promega) were incubated with His-OsMKK1/OsMKK6 in GST binding buffer (2 mM KH_2_PO_4_, 4.2 mM Na_2_HPO_4_, 10 mM KCl, 140 mM NaCl, 10% bovine serum albumin, pH 7.5) at 4°C for 1 h. The particles were washed three times with GST wash buffer (similar with GST binding buffer but without bovine serum albumin), separated on 12% SDS-PAGE gel and detected by immunoblotting with anti-His antibody (Abmart, lot: 283874, 1:1000, v/v) or anti-GST antibody (Abmart, lot:264160; 1:1000, v/v).

### BiFC Assay

Full-length coding regions of *OsMKK1*/*OsMKK6* were separately amplified and cloned into the *BamH*I-*Kpn*I sites and the *Cla*I-*Kpn*I sites of the pSPYNE vector to generate the *pSPYNE*-*OsMKK1/OsMKK6* constructs. The *pSPYCE*-*OsDMI3* construct was generated as described previously (Ni et al., 2019). The corresponding constructs were transiently co-transformed in onion epidermis cells using the particle bombardment (Bio-Rad) method as described previously (Lee *et al*., 2008). The YFP fluorescent signals were observed at 16 h after transformation using a confocal scanning microscope (Zeiss, German).

### Co-IP Assay

For Co-IP assay, the construct *OsDMI3-Myc* was generated as described previously (Ni et al., 2019). *OsMKK1*/*OsMKK6* were fused with *Flag* and separately cloned into the pXZP008 vector at *BamH*I-*Kpn*I sites and *SaI*I-*Kpn*I sites to generate *OsMKK1*/*OsMKK6-Flag* constructs. The constructs were transiently expressed in rice protoplasts via PEG-mediated transfection method (Ni et al., 2019). After 16 h incubation, the proteins of transfected protoplasts were extracted with Co-IP buffer (20 mM Tris-HCl, pH 7.5, 150 mM NaCl, 1 mM EDTA, 10 mM Na_3_VO_4_, 10 mM NaF, 10% glycerol, 5 μg mL^-1^ leupeptin, 5 μg mL^-1^ aprotinin, 0.5% (v/v) Triton X-100, 0.5% (v/v) Nonidet P-40). The proteins were incubated with anti-Myc antibody bound to protein A beads for 2 to 3 h. The beads were collected and washed three times with wash buffer (20 mM Tris-HCl, pH 7.5, 150 mM NaCl, 1 mM EDTA, 10 mM Na_3_VO_4_, 10 mM NaF, 10% glycerol, 0.5% [v/v] Triton X-100, 0.5% [v/v] Nonidet P-40), and then were boiled in 1×SDS loading buffer. The immunoprecipitated proteins were analyzed by SDS-PAGE and immunoblotting with anti-Flag antibody (Abmart, lot:293674; 1:1000, v/v).

### LCI Assay

For LCI assay, the full-length *OsMKK1* and *OsDMI3* were ligated into the *BamH*I/*Kpn*I sites of the pC1300-nLUC vector and the *Kpn*I*/Pst*I sites of the pC1300-cLUC vector, respectively. Both the nLUC- and cLUC-fused constructs were transformed into *Nicotiana benthamiana* leaves using the *Agrobacterium*-mediated infiltration method (Ni et al., 2019). After 3 days of infiltration, luciferase activity was analyzed using chemiluminescence imaging (Tanon 5200 Multi, Tanon Biomart).

### Immunocomplex Kinase Activity Assay

Protein was extracted from rice leaves with buffer (100 mM Hepes, 10 mM Na_3_VO_4_, 10 mM NaF, 10% glycerol, 5 μg mL^-1^ leupeptin, 5 μg mL^-1^ aprotinin, 1 mM PMSF, 10 mM DTT). Protein content was determined by the method of Bradford (1976). For the determination of OsMKK1 and OsMKK6 activity, the total proteins (200 μg) were bound with anti-OsMKK1 antibody or anti-OsMKK6 antibody (2 μg, Abmart) in immunoprecipitation buffer (20 mM Tris-HCl, pH 7.5, 150 mM NaCl, 1 mM EDTA, 10 mM Na_3_VO_4_, 10 mM NaF, 10% glycerol, 0.5% [v/v] Triton X-100, 0.5% [v/v] Nonidet P-40) overnight, and then incubated with protein A-sepharose beads for another 3 h. The immunoprecipitated proteins were incubated with 1 μg MBP (Sigma-Aldrich) in kinase buffer (25 mM Tris-HCl, pH 7.5, 5 mM MgCl_2_, 1 mM ethylene glycol tetraacetic acid [EGTA], 0.1 mM Na_3_VO_4_ and 1 mM DTT) contained 10 μCi [γ^32^P]-ATP (3000 Ci mM^-1^) at 30°C for 30 min. The reaction products were separated by 12% SDS-PAGE and analyzed by autoradiography using X-ray film or a phosphostorage screen (Typhoon TRIO, Amersham Biosciences). For the determination of OsDMI3 and OsMPK1 activity, the immunocomplex kinase activity assay was performed as described previously (Ni et al., 2019). The relative levels of OsDMI3, OsMKK1, OsMKK6, and OsMPK1 activity were quantified by ImageJ, and were presented as values relative to those of the corresponding controls.

### SDS-PAGE and Immunoblot Analysis

Protein extracts (20 μg) were separated by 12% SDS-PAGE. After electrophoresis, the gel was transferred to a polyvinylidene difluoride (PVDF) membrane, and then incubated in phosphate buffered saline/Tween (PBST) buffer (140 mM NaCl, 10 mM KCl, 2 mM KH_2_PO_4_, 8 mM Na_2_HPO_4_, 0.1% Tween-20 [v/w], pH 7.5) contained 5% (w/v) nonfat dry milk for 2 h at room temperature. The membrane was then washed three times with PBST buffer for 5 min. The blots were probed with anti-OsDMI3 antibody (ABclonal), anti-OsMKK1 antibody (Abmart), anti-OsMKK6 antibody (Abmart), anti-OsMPK1 antibody (Beijing Protein Innovation), anti-pT25 OsMKK1 antibody (GenScript), anti-ACT1 antibody (Beijing Protein Innovation), anti-Myc antibody, anti-Flag antibody, anti-His antibody, and anti-GST antibody. The information about anti-OsDMI3 antibody was described previously (Shi et al., 2012). The anti-OsMKK1 antibody was raised against peptide (KDGLRIVSQSEE) of OsMKK1, the anti-OsMKK6 antibody was raised against peptide (IKKFEDKDLDLR) of OsMKK6, and the anti-pT25 OsMKK1 antibody was raised against peptide (LTQSG**T**FKDGDLLVN) of OsMKK1. First antibody and secondary antibody were used at 1:1000 and 1:5000 dilution, respectively. Chemiluminescence was detected with the enhanced chemiluminescence immunoblotting detection system (GE Healthcare) and a camera (Tanon 5200 Multi, Tanon).

### In Vitro Kinase Assay

Total protein of rice leaves was extracted as described above, and the protein extract (200 μg) was incubated with anti-OsDMI3 antibody (2 μg, Abmart) and immunoprecipitation buffer (20 mM Tris-HCl, pH 7.5, 150 mM NaCl, 1 mM EDTA, 10 mM Na_3_VO_4_, 10 mM NaF, 10% glycerol, 0.5% [v/v] Triton X-100, 0.5% [v/v] Nonidet P-40) to obtain the kinase OsDMI3. OsDMI3 was then incubated with 20 μg substrates (His-OsMPK1, His-OsMKK6, His-OsMKK1, and His-OsMKK1^T25A^) and 10 μCi [γ^32^P]-ATP in reaction buffer (25 mM Tris-HCl, pH 7.5, 5 mM MgCl_2_, 0.5 mM CaCl_2_, 2 μM CaM [Sigma-Aldrich], 1 mM DTT) at 30°C for 30 min to perform the vitro kinase assay. The reaction was stopped by mixing SDS loading buffer, and the reaction mixtures were separated by 12% SDS-PAGE. The phosphorylated substrates were visualized by autoradiography.

For the in vitro kinase assay of OsMKK1 activity, 10 μg of His-OsMKK1, His-OsMKK1^T25A^, His-OsMKK1^T25D^, His-OsMKK1^S215A/T221A^, and His-OsMKK1^S215D/T221D^ were respectively incubated with 1 μg MBP in kinase buffer (25 mM Tris-HCl, pH 7.5, 5 mM MgCl_2_, 1 mM EGTA, 0.1 mM Na_3_VO_4_ and 1 mM DTT) and 10 µCi [γ-^32^P]-ATP at 30°C for 30 min. SDS sample buffer was then added to stop the reaction. After separation by 12% SDS-PAGE, the phosphorylated MBP was visualized by autoradiography.

### In Vitro Reconstitution Assay of OsMPK1 phosphorylation

10 μg of His-OsMKK1 or His-OsMKK1^T25A^ or His-OsMKK1^T25D^ was incubated with 20 μg GST-OsMPK1 in reaction buffer (25 mM Tris-HCl, pH 7.5, 5 mM MgCl_2_, 2.5 mM MnCl_2,_ 0.5 mM CaCl_2_ and 1 mM DTT) contained 10 µCi [γ-^32^P]-ATP at 30°C for 30 min. The reaction mixtures were then incubated with or without OsDMI3 pulled down from extracts of ABA-treated leaves at 30°C for another 30 min. The phosphorylated OsMPK1 was analyzed as described in vitro kinase assay.

### In Vivo Phosphorylation Assay

OsMKK1 and OsMKK6 were expressed in rice protoplasts from WT and *osdmi3*-KO1 as Flag-tagged proteins, and the total proteins in the protoplasts after treatment with 10 μM ABA for 5, 10, and 15 min were extracted in buffer (20 mM Tris-HCl, pH 7.5, 150 mM NaCl, 1 mM EDTA, 10 mM Na_3_VO_4_, 10 mM NaF, 10% glycerol, 5 μg mL^-1^ leupeptin, 5 μg mL^-1^ aprotinin, 0.5% [v/v] Triton X-100, 0.5% [v/v] Nonidet P-40, 1% [v/v] phosphatase inhibitor cocktail 3 [Sigma-Aldrich]). OsMKK1-Flag and OsMKK6-Flag proteins were immunoprecipitated by anti-Flag antibody bound to protein A beads, and separated by 12% SDS-PAGE. Phosphorylated proteins were detected by immunoblotting using Biotinylated Phos-tag as described previously (Kinoshita-Kikuta et al., 2007).

### Mass Spectrometry Analysis

OsMKK1 was incubated with 1 μg calf intestine alkaline phosphatase (CIAP) in 100 μL of phosphatase buffer (50 mM KAc, 20 mM Tris-HAc, pH 7.9, 10 mM Mg(Ac)_2_, 100 μg mL^-1^ BSA) at 37°C for 10 min. The phosphatase was deactivated by heating at 80°C for 2 min. The dephosphorylated OsMKK1 was then incubated with OsDMI3 and ATP (200 nM) in kinase reaction solution (25 mM Tris-HCl, pH 7.5, 5 mM MgCl_2_, 0.5 mM CaCl_2_, 2 μM CaM, 1 mM DTT) at 30°C for 30 min. The phosphorylated OsMKK1 was digested by trypsin and analyzed by mass spectrometry as described previously (Gampala et al., 2007). Phosphopeptide sequence of OsMKK1 identified by LC-MS/MS analysis is listed in Supplemental Table 1.

### Parallel Reaction Monitoring (PRM) Analysis

*OsMKK1-Flag* was introduced into rice protoplasts of *OsDMI3*-OE1, *osdmi3*-KO1 and WT respectively, and the transfected protoplasts were treated with 10 μM ABA for 10 min after 16 h incubation. Protein was extracted from protoplasts with buffer (20 mM Tris-HCl, pH 7.5, 150 mM NaCl, 1 mM EDTA, 10 mM Na_3_VO_4_, 10 mM NaF, 10% glycerol, 5 μg mL^-1^ leupeptin, 5 μg mL^-1^ aprotinin, 0.5% [v/v] Triton X-100, 0.5% [v/v] Nonidet P-40, 1% [v/v] phosphatase inhibitor cocktail 3 [Sigma-Aldrich]), and the protein extracts were immunoprecipitated using anti-Flag antibody and separated by 12% SDS-PAGE. A pre-test was performed to verify if the target phosphopeptides were detected by mass spectrometry after the OsMKK1 protein was digested by trypsin. LC-PRM MS analysis (Peterson et al., 2012) was applied to quantify the target phosphopeptides. Briefly, the label free protocol was used for phosphopeptides 2 preparation. Each sample was spiked with the stable isotope-containing AQUA peptide as a standard internal reference, and tryptic peptides were loaded on stage tips of C18 for desalting prior to reversed-phase chromatography on one of the nLC-1200 easy systems (Thermo Scientific). Then, 1 h liquid chromatography gradients were performed with 5-35% acetonitrile for 45 min. After these, Q Exactive Plus MS was applied for PRM analysis. The analysis of raw data was realized via Skyline (MacCoss Lab, University of Washington; MacLean et al., 2010), wherein the intensity of signal produced by a certain phosphopeptides sequence could be quantified with respect to each sample and referenced to standards via normalization for each protein.

### Site-Directed Mutagenesis

The mutated variants were achieved by site-directed mutagenesis using the Multi-Directed Mutagenesis Kit (Agilent Technologies: California) according to the manufacturer’s instructions. The sequences of DNA oligonucleotides used in mutagenesis are listed in Supplemental Table 2.

### Quantitative RT-PCR Analyses

Total RNA was extracted with RNAiso Plus kit (TaKaRa). Approximately 2 μg of total RNA was reverse-transcribed using PrimeScript™ RT reagent Kit with gDNA Eraser (TaKaRa). Real-time quantitative RT-PCR was performed with TB Green Premix Ex Taq™ (TakaRa) using a 7500 real-time PCR system (Applied Biosystems Inc.). The expression level was normalized against that of rice *glyceraldehyde-3-phosphate dehydrogenase* (*GAPDH*) gene. The primers used are listed at Supplemental Table 3.

### Phenotype Analysis

For seed germination assay, the seeds of WT, *OsMKK1, osmkk1, OsMKK1*^*T25A*^, *OsMKK1*^*T25D*^, *OsDMI3, osdmi3, osmkk1*/*osdmi3, osmkk1*/*OsDMI3, osdmi3*/*OsMKK1, osdmi3*/*OsMKK1*^*T25A*^, and *osdmi3*/*OsMKK1*^*T25D*^ were surface-sterilized and germinated on half-strength Murashige and Skoog medium (0.8% [w/v] agar) with different concentrations of ABA (0, 1, and 5 μM). The seeds were incubated at 4°C for 2 d and then transferred to a growth chamber (16 h light/8 h dark; 200 μmol m^−2^ s^−1^ light intensity; 25°C) to germinate. The germination rate of seeds was scored at the indicated times.

For the detection of root growth, 4-d-old rice seedlings grown in the normal conditions were transferred to a nutrient solution containing different concentrations of ABA (0, 0.5, 1, 2.5, 5, and 10 μM) for another 8 d, and then the length of primary roots was measured.

For the analysis of survival rate, 10-d-old rice seedings were treated with 20% PEG 4000, 100 mM H_2_O_2_ for 12 d or 18 d and the survival rates were measured after recovery by re-watering for 7 d.

For the determination of oxidative damage to lipids and plasma membranes, the rice seedings were treated with 20% PEG 4000 or 100 mM H_2_O_2_ for 2 d, and the content of MDA and the percentage of electrolyte leakage were determined by the methods described previously (Jiang and Zhang, 2001).

For the determination of water loss, the fully expanded leaves of 10-d-old rice seedlings were detached, weighed, and placed on the laboratory bench. Weight loss of the detached leaves was monitored at the indicated time intervals as shown in Supplemental Figure 17C. Water loss was expressed as the percentage of initial fresh weight.

### Determination of ABA Content

Fresh leaves (0.5 g) of 10-d-old rice seedlings were collected, frozen in liquid nitrogen and ground in a mortar, and extracted in 7 mL extraction solution (methanol:water:acetic acid, 80:19:1) overnight at 4°C. After centrifugation at 8000 g for 20 min, the supernatant was collected and eluted through a Sep-Pak C18 cartridge (Waters, Milford, MA, USA), which was preconditioned with methanol. The total extract solution was dried under nitrogen gas and then dissolved in 500 μL mobile phase (methanol:1% acetic acid, 45: 55). The extract was filtered through 0.45 μm membrane filters before injection into the high-performance liquid chromatography (HPLC). Detection was done with an absorbance detector at a wavelength of 254 nm. Quantification was obtained by comparing the peak areas with those of known amounts of ABA (Sigma, Shanghai, China).

### Alignment Analysis

Sequences of plant MKK1 proteins from different species were retrieved from NCBI and aligned by Alignx. The GenBank accession numbers are as follows: *ZmMKK1*, BT065734; *AtMKK1*, AT4g26070; *GmMKK1*, AY070230; *MtMKK1*, AC144503.

### Statistical Analysis

Statistical analysis was conducted using the program SPSS Statistics v22.0 (IBM). Data were analyzed by ANOVA followed by the Duncan’s multiple range test (P < 0.05 as the level of significance).

### Accession Numbers

Sequence data from this article can be found in the GenBank/EMBL data libraries under the following accession numbers: *OsDMI3*, LOC_Os05g41090; *OsMPK1*, LOC_Os06g06090; *OsACT1*, LOC_Os03g50885; *OsMKK1*, LOC_Os06g05520; *OsMKK6*, LOC_Os01g32660; *OsMKK3*, LOC_Os06g27890; *OsMKK4*, LOC_Os02g54600; *OsMKK5*, LOC_Os06g09180; *OsMKK10-2*, LOC_Os03g12390; *OsABA2*, LOC_Os03g59610; *OsGAPDH*, LOC_Os02g38920.

## Supplemental Data

**Supplemental Figure 1**. OsDMI3 Does Not Interact with OsMPK1 and Does Not Phosphorylate OsMPK1.

**Supplement Figure 2**. OsDMI3 Interacts with the Group A MKKs, OsMKK1 and OsMKK6.

**Supplemental Figure 3**. OsMKK1 Interacts with the EF Hands Domain of OsDMI3.

**Supplemental Figure 4**. OsDMI3 Interacts with the N-Terminus Domain of OsMKK1.

**Supplemental Figure 5**. OsMKK6 Interacts with the EF Hands Domain of OsDMI3.

**Supplemental Figure 6**. OsDMI3 Interacts with the N-Terminus Domain of OsMKK6.

**Supplemental Figure 7**. Identification of *osmkk1*-KO and *osmkk6*-KO Mutants.

**Supplement Figure 8**. The Specificity of the Anti-OsMKK1 Antibody and the Anti-OsMKK6 Antibody.

**Supplemental Figure 9**. Ca^2+^ Is Required for ABA-Induced Activation of OsMKK1.

**Supplemental Figure 10**. ABA-Induced Activation of OsMKK6 Is Ca^2+^-Independent.

**Supplemental Figure 11**. OsMKK1 Does Not Affect the Activation of OsDMI3 in ABA signaling.

**Supplemental Figure 12**. Identification of the Phosphorylation Site of OsMKK1 by OsDMI3 In Vitro.

**Supplemental Figure 13**. The Specificity of the Anti-pT25 OsMKK1 Antibody.

**Supplemental Figure 14**. Identification of *osaba2*-KO Mutant.

**Supplemental Figure 15**. The Expression of *OsMKK1* and *OsDMI3* and the Levels of OsMKK1 and OsDMI3 Proteins in the Transgenic Lines.

**Supplemental Figure 16**. Phosphorylation of OsMKK1 at Thr-25 Does Not Affect the Interaction between OsDMI3 with OsMKK1 and between OsMKK1 with OsMPK1.

**Supplemental Figure 17**. The OsDMI3-OsMKK1 Pathway Regulates the Physiological Response to Water Stress and Oxidative Stress.

**Supplemental Table 1**. Phosphopetide Sequence of OsMKK1 Identified by LC-MS/MS Analysis.

**Supplemental Table 2**. DNA Oligonucleotides Used for Site-Directed Mutagenesis.

**Supplemental Table 3**. PCR Primers Used in This Study.

## ACKNOWLEDGMENTS

This work was supported by the National Natural Science Foundation of China (grant no. 31671606 and 31971824) and the National Basic Research Program of China (grant no. 2012CB114306).

## AUTHOR CONTRIBUTIONS

M. J. conceived the project. M.J. and M.C. designed the experiments. M.C. performed most of the experiments. J.C., M.S., C.Q., and G.Z. performed some of the experiments. M.J., L.N., and A.Z. analyzed data. M.J., M.C., and L.N. wrote the manuscript.

## Parsed Citations

Bai, F., and Matton, D.P. (2018). The Arabidopsis Mitogen-Activated Protein Kinase Kinase Kinase 20 (MKKK20) C-terminal domain interacts with MKK3 and harbors a typical DEF mammalian MAP kinase docking site. Plant Signal. Behav. 13: e1503498

Bardwell, L. (2006). Mechanisms of MAPK signaling specificity. Biochem. Soc. Trans. 34: 837–841.

Bi, G., Zhou, Z., Wang, W., Li, L., Rao, S., Wu, Y., Zhang, X., Menke, F.L.H., Chen, S., and Zhou, J.M. (2018). Receptor-like cytoplasmic kinases directly link diverse pattern recognition receptors to the activation of mitogen-activated protein kinase cascades in Arabidopsis. Plant Cell 30: 1543–1561.

Bigeard, J., and Hirt, H. (2018). Nuclear signaling of plant MAPKs. Front. Plant Sci. 9: 469.

Bradford, M.M. (1976). Arapid and sensitive method for the quantitation of microgram quantities of protein utilizing the principle of protein–dye binding.Anal. Biochem. 72: 248–254.

Cheng, W.H., et al. (2002). Aunique short-chain dehydrogenase/reductase in Arabidopsis glucose signaling and abscisic acid biosynthesis and functions. Plant Cell 14: 2723–2743.

Colcombet, J., and Hirt, H. (2008). Arabidopsis MAPKs: a complex signaling network involved in multiple biological processes. Biochem. J. 413: 217–226.

Cutler, S.R., Rodriguez, P.L., Finkelstein, R.R., and Abrams, S.R. (2010) Abscisic acid: emergence of a core signaling network. Annu. Rev. Plant Biol. 61: 651–679.

Danquah, A., et al. (2015). Identification and characterization of an ABA-activated MAP kinase cascade in Arabidopsis thaliana. Plant J. 82: 232–244.

Danquah, A., de Zélicourt, A., Colcombet, J., and Hirt, H. (2014). The role of ABA and MAPK signaling pathways in plant abiotic stress responses. Biotechnol. Adv. 32: 40–52.

de Zelicourt, A., Colcombet, J., and Hirt, H. (2016). The role of MAPK modules and ABA during abiotic stress signaling. Trends Plant Sci. 21: 677–685.

Dóczi, R., and Bögre, L. (2018). The quest for MAP kinase substrates: gaining momentum. Trends Plant Sci. 23: 918–932.

Furuya, T., Matsuoka, D., and Nanmori, T. (2013). Phosphorylation of Arabidopsis thaliana MEKK1 via Ca2+ signaling as a part of the cold stress response. J. Plant Res. 126: 833–840.

Gampala, S.S., et al. (2007). An essential role for 14-3-3 proteins in brassinosteroid signal transduction in Arabidopsis. Dev. Cell 13: 177–189.

Gao, M., Liu, J., Bi, D., Zhang, Z., Cheng, F., Chen, S., and Zhang, Y. (2008). MEKK1, MKK1/MKK2 and MPK4 function together in a mitogen-activated protein kinase cascade to regulate innate immunity in plants. Cell Res. 18: 1190–1198.

Ge, B., Gram, H., Di Padova, F., Huang, B., New, L., Ulevitch, R.J., Luo, Y., and Han, J. (2002). MAPKK-independent activation of p38(mediated by TAB1-dependent autophosphorylation of p38(. Science 295: 1291–1294.

González-Guzmán, M., Apostolova, N., Bellés, J.M., Barrero, J.M., Piqueras, P., Ponce, M.R., Micol, J.L., Serrano, R., and Rodríguez, P.L. (2002). The short-chain alcohol dehydrogenase ABA2 catalyzes the conversion of xanthoxin to abscisic aldehyde. Plant Cell 14: 1833–1846.

Hamel, L.P., Nicole, M.C., Duplessis, S., and Ellis, B.E. (2012). Mitogen-activated protein kinase signaling in plant-interacting fungi: distinct messages from conserved messengers. Plant Cell 24: 1327–1351.

Hamel, L.P., et al. (2006). Ancient signals: comparative genomics of plant MAPK and MAPKK gene families. Trends Plant Sci. 11: 192–198.

Huang, R., Zheng, R., He, J., Zhou, Z., Wang, J., Xiong, Y., and Xu, T. (2019). Noncanonical auxin signaling regulates cell division pattern during lateral root development. Proc. Natl. Acad. Sci. USA 116: 21285–21290.

Jagodzik, P., Tajdel-Zielinska, M., Ciesla, A., Marczak, M., and Ludwikow, A. (2018). Mitogen-activated protein kinase cascades in plant hormone signaling. Front. Plant Sci. 9: 1387.

Jiang, M., and Zhang, J. (2001). Effect of abscisic acid on active oxygen species, antioxidative defence system and oxidative damage in leaves of maize seedlings. Plant Cell Physiol. 42: 1265–1273.

Kinoshita-Kikuta, E., Aoki, Y., Kinoshita, E., and Koike, T. (2007). Label-free kinase profiling using phosphate affinity polyacrylamide gel electrophoresis. Mol. Cell. Proteomics 6: 356–366.

Krysan, P.J., and Colcombet, J. (2018). Cellular complexity in MAPK signaling in plants: questions and emerging tools to answer them. Front. Plant Sci. 9: 1674.

Lee, M.O., et al. (2008). Novel rice OsSIPK is a multiple stress responsive MAPK family member showing rhythmic expression at mRNA level. Planta 227: 981–990.

Li, K., Yang, F., Zhang, G., Song, S., Li, Y., Ren, D., Miao, Y., and Song, C.P. (2017). AIK1, A mitogen-activated protein kinase, modulates abscisic acid responses through the MKK5-MPK6 kinase cascade. Plant Physiol. 173: 1391–1408.

Ma, F., Lu, R., Liu, H., Shi, B., Zhang, J., Tan, M., Zhang, A., and Jiang, M. (2012). Nitric oxide-activated calcium/calmodulin-dependent protein kinase regulates the abscisic acid-induced antioxidant defence in maize. J. Exp. Bot. 63: 4835–4847.

Ma, F., Ni, L., Liu, L., Li, X., Zhang, H., Zhang, A., Tan, M., and Jiang, M. (2016). ZmABA2, an interacting protein of ZmMPK5, is involved in abscisic acid biosynthesis and functions. Plant Biotechnol. J. 14: 771–782.

MacLean, B., Tomazela, D.M., Shulman, N., Chambers, M., Finney, G.L., Frewen, B., Kern, R., Tabb, D.L., Liebler, D.C., MacCoss, M.J. (2010). Skyline: an open source document editor for creating and analyzing targeted proteomics experiments. Bioinformatics 26: 966–968.

MAPK Group. (2002). Mitogen-activated protein kinase cascades in plants: a new nomenclature. Trends Plant Sci. 7: 301–308.

Matsuoka, D., Yasufuku, T., Furuya, T., and Nanmori, T. (2015). An abscisic acid inducible Arabidopsis MAPKKK, MAPKKK18 regulates leaf senescence via its kinase activity. Plant Mol. Biol. 87: 565–575.

Miller, C.J., and Turk, B.E. (2018). Homing in: mechanisms of substrate targeting by protein kinases. Trends Biochem. Sci. 43: 380–394.

Ni, L., et al. (2019). Abscisic acid inhibits rice protein phosphatase PP45 via H2O2 and relieves repression of the Ca2+/CaM-dependent protein kinase DMI3. Plant Cell 31: 128–152.

Peterson, A.C., Russell, J.D., Bailey, D.J., Westphall, M.S., and Coon, J.J. (2012). Parallel reaction monitoring for high resolution and high mass accuracy quantitative, targeted proteomics. Mol. Cell. Proteomics 11: 1475–1488.

Pitzschke, A. (2015). Modes of MAPK substrate recognition and control. Trends Plant Sci. 20: 49–55.

Poovaiah, B.W., Du, L., Wang, H., and Yang, T. (2013). Recent advances in calcium/ calmodulin-mediated signaling with an emphasis on plant-microbe interactions. Plant Physiol. 163: 531–542.

Rao, K.P., Richa, T., Kumar, K., Raghuram, B., and Sinha, A.K. (2010). In silico analysis reveals 75 members of mitogen-activated protein kinase kinase kinase gene family in rice. DNARes. 17: 139–153.

Ren, D., Yang, H., and Zhang, S. (2002). Cell death mediated by MAPK is associated with hydrogen peroxide production in Arabidopsis. J. Biol. Chem. 277: 559–565.

Rodriguez, M.C., Petersen, M., and Mundy, J. (2010). Mitogen-activated protein kinase signaling in plants. Annu. Rev. Plant Biol. 61: 621–649.

Salvador, J.M., Mittelstadt, P.R., Guszczynski, T., Copeland, T.D., Yamaguchi, H., Appella, E., Fornace Jr, A.J., and Ashwell, J.D. (2005). Alternative p38 activation pathway mediated by T cell receptor-proximal tyrosine kinases. Nat. Immunol. 6: 390–395.

Šamajová, O., Plíhal, O., Al-Yousif, M., Hirt, H., and Šamaj, J. (2013). Improvement of stress tolerance in plants by genetic manipulation of mitogen-activated protein kinases. Biotechnol. Adv. 31: 118–128.

Shi, B., Ni, L., Liu, Y., Zhang, A., Tan, M., and Jiang, M. (2014). OsDMI3-mediated activation of OsMPK1 regulates the activities of antioxidant enzymes in abscisic acid signaling in rice. Plant Cell Environ. 37: 341–352.

Shi, B., Ni, L., Zhang, A., Cao, J., Zhang, H., Qin, T., Tan, M., Zhang, J., and Jiang, M. (2012). OsDMI3 is a novel component of abscisic acid signaling in the induction of antioxidant defense in leaves of rice. Mol. Plant 5: 1359–1374.

Singh, R., et al. (2012). Rice mitogen-activated protein kinase interactome analysis using the yeast two-hybrid system. Plant Physiol. 160: 477–487.

Singh, R., and Jwa, N.S. (2013). The rice MAPKK–MAPK interactome: the biological significance of MAPK components in hormone signal transduction. Plant Cell Rep. 32: 923–931.

Singh, S., and Parniske, M. (2012). Activation of calcium-and calmodulin-dependent protein kinase (CCaMK), the central regulator of plant root endosymbiosis. Curr. Opin. Plant Biol. 15: 444–453.

Suarez-Rodriguez, M.C., Adams-Phillips, L., Liu, Y., Wang, H., Su, S.H., Jester, P.J., Zhang, S., Bent, A.F., and Krysan, P.J. (2007). MEKK1 is required for flg22-induced MPK4 activation in Arabidopsis plants. Plant Physiol. 143: 661–669.

Sun, T., Nitta, Y., Zhang, Q., Wu, D., Tian, H., Lee, J.S., and Zhang, Y. (2018). Antagonistic interactions between two MAP kinase cascades in plant development and immune signaling. EMBO Rep. 19: 45324.

Takahashi, F., Mizoguchi, T., Yoshida, R., Ichimura, K., and Shinozaki, K. (2011). Calmodulin-dependent activation of MAP kinase for ROS homeostasis in Arabidopsis.Mol. Cell 41: 649–660.

Takekawa, M., Tatebayashi, K., and Saito, H. (2005). Conserved docking site is essential for activation of mammalian MAP kinase kinases by specific MAP kinase kinase kinases. Mol. Cell 18: 295–306.

Ubersax, J.A., and Ferrell, J.E. (2007). Mechanisms of specificity in protein phosphorylation. Nat. Rev. Mol. Cell Biol. 8: 530–541.

Umezawa, T., Nakashima, K., Miyakawa, T., Kuromori, T., Tanokura, M., Shinozaki, K., and Yamaguchi-Shinozaki, K. (2010). Molecular basis of the core regulatory network in ABA responses: sensing, signaling and transport. Plant Cell Physiol. 51: 1821–1839.

Umezawa, T., Takahashi, F., and Shinozaki, K. (2014). Phosphorylation networks in the abscisic acid signaling pathway. Enzymes 35: 27–56.

Wang, C., Wang, G., Zhang, C., Zhu, P., Dai, H., Yu, N., He, Z., Xu, L., and Wang, E. (2017). OsCERK1-mediated chitin perception and immune signaling requires receptor-like cytoplasmic kinase 185 to activate an MAPK cascade in rice. Mol. Plant 10: 619–633.

Wang, J.P., Munyampundu, J.P., Xu, Y.P., and Cai, X.Z. (2015). Phylogeny of plant calcium and calmodulin-dependent protein kinases (CCaMKs) and functional analyses of tomato CCaMK in disease resistance. Front. Plant Sci. 6: 1075.

Xie, K., Chen, J., Wang, Q., and Yang, Y. (2014). Direct phosphorylation and activation of a mitogen-activated protein kinase by a calcium-dependent protein kinase in rice. Plant Cell 26: 3077–3089.

Xing, Y., Jia, W., and Zhang, J. (2008). AtMKK1 mediates ABA-induced CAT1 expression and H2O2 production via AtMPK6-coupled signaling in Arabidopsis. Plant J. 54: 440–451.

Xu, J., and Zhang, S. (2015). Mitogen-activated protein kinase cascades in signaling plant growth and development. Trends Plant Sci. 20: 56–64.

Yamada, K., Yamaguchi, K., Yoshimura, S., Terauchi, A., and Kawasaki, T. (2017). Conservation of chitin-induced MAPK signaling pathways in rice and Arabidopsis. Plant Cell Physiol. 58: 993–1002.

Yang, T., Chaudhuri, S., Yang, L., Du, L., and Poovaiah, B.W. (2010a). Acalcium/calmodulin-regulated member of the receptor-like kinase family confers cold tolerance in plants. J. Biol. Chem. 285: 7119–7126.

Yang, T., Shad Ali, G., Yang, L., Du, L., Reddy, A.S.N., and Poovaiah, B.W. (2010b). Calcium/calmodulin-regulated receptor-like kinase CRLK1 interacts with MEKK1 in plants. Plant Signal. Behav. 5: 991–994.

Yu, L., Nie, J., Cao, C., Jin, Y., Yan, M., Wang, F., Liu, J., Xiao, Y., Liang, Y., and Zhang, W. (2010). Phosphatidic acid mediates salt stress response by regulation of MPK6 in Arabidopsis thaliana. New Phytol. 188: 762–773.

Zhang, H., Liu, Y., Wen, F., Yao, D., Wang, L., Guo, J., Ni, L., Zhang, A., Tan, M., and Jiang, M. (2014). Anovel rice C2H2-type zinc finger protein, ZFP36, is a key player involved in abscisic acid-induced antioxidant defence and oxidative stress tolerance in rice. J. Exp. Bot. 65: 5795–5809.

Zhang, M., Su, J., Zhang, Y., Xu, J., and Zhang, S. (2018). Conveying endogenous and exogenous signals: MAPK cascades in plant growth and defense. Curr. Opin. Plant Biol. 45: 1–10.

Zhang, Z., Ke, D., Hu, M., Zhang, C., Deng, L., Li, Y., Li, J., Zhao, H., Cheng, L., Wang, L., and Yuan, H. (2019). Quantitative phosphoproteomic analyses provide evidence for extensive phosphorylation of regulatory proteins in the rhizobia–legume symbiosis. Plant Mol. Biol. 100: 265–283.

Zhao, C., Wang, P., Si, T., Hsu, C.C., Wang, L., Zayed, O., Yu, Z., Zhu, Y., Dong, J., Tao, W.A., and Zhu, J.K. (2017). MAP kinase cascades regulate the cold response by modulating ICE1 protein stability. Dev. Cell 43: 618–629.

Zhu, J.K. (2016a). Abiotic stress signaling and responses in plants. Cell 167: 313–324.

Zhu, Y., et al. (2016b). Phosphorylation of a NAC transcription factor by ZmCCaMK regulates abscisic acid-induced antioxidant defense in maize. Plant Physiol. 171: 1651–1664.

